# Megaphage infect *Prevotella* and variants are widespread in gut microbiomes

**DOI:** 10.1101/356790

**Authors:** Audra E. Devoto, Joanne M. Santini, Matthew R. Olm, Karthik Anantharaman, Patrick Munk, Jenny Tung, Elizabeth A. Archie, Peter J. Turnbaugh, Kimberley D. Seed, Ran Blekhman, Frank M. Aarestrup, Brian C. Thomas, Jillian F. Banfield

## Abstract

Bacteriophage (phage) dramatically shape microbial community composition, redistribute nutrients *via* host lysis, and drive evolution through horizontal gene transfer. Despite their importance, much remains to be learned about phage in the human microbiome. We investigated gut microbiomes of humans from Bangladesh and Tanzania, two African baboon social groups, and Danish pigs, and report that many contain phage belonging to a clade with genomes >540 kb in length, the largest yet reported in the human microbiome and close to the maximum size ever reported for phage. We refer to these as Lak phage. CRISPR spacer targeting indicates that the Lak phage infect bacteria of the genus *Prevotella*. We manually curated to completion 15 distinct Lak phage genomes recovered from metagenomes. The genomes display several interesting features, including use of an alternative genetic code, large intergenic regions that are highly expressed, and up to 35 putative tRNAs, some of which contain enigmatic introns. Different individuals have distinct phage genotypes, and shifts in variant frequencies over consecutive sampling days reflect changes in relative abundance of phage sub-populations. Recent homologous recombination has resulted in extensive genome admixture of nine baboon Lak phage populations. We infer that Lak phage are widespread in gut communities that contain *Prevotella* species, especially in individuals in the developing world, and conclude that megaphage, with fascinating and underexplored biology, may be common but largely overlooked components of human and animal gut microbiomes.

## INTRODUCTION

The compositions and dynamics of human and animal microbiomes are of enormous interest, given that microbial activity impacts nutrition, physiological development, and disease (1). The human gut microbiome has been intensively studied, mostly using gene fingerprinting methods to resolve body site specificity and microbiome compositional changes with age, health conditions and diet (2–4). Less commonly applied is genome-resolved metagenomics, which involves simultaneous recovery of draft and sometimes complete genomes from metagenomes. Such studies have genomically described novel bacterial and archaeal species and genera and in one case, a new phylum (5),and provided access to bacteriophage (phage), virus and plasmid sequences that are not accessible *via* fingerprinting methods (6–8).

Phage are increasingly recognized as ubiquitous components of microbiomes. They can dramatically shape ecosystem structure *via* strain-specific predation, mediate horizontal gene transfer, and redistribute nutrients by lysing host cells (9). Recently, crAssphage were discovered in human microbiomes (10). The ~97 kbp sequence is highly represented in public metagenomes and accounted for 1.68% of human faecal metagenomic sequencing reads. Viral particles closely related to crAssphage were recently amplified *ex vivo*, but they have not yet been isolated (11). As the crAssphage host is predicted to be *Bacteroides*, crAssphage may often be associated with people colonized by *Bacteroides*, possibly due to consumption of a Western diet (12).

We began the current study by investigating microbial communities in the gastrointestinal tracts of ten arsenic-impacted adult men from Laksam Upazila, Bangladesh. Our objectives were to identify new gut microbiome-associated phages, link them to bacterial hosts and evaluate their distribution. In faecal samples from these individuals we discovered phage with genomes that are exceptionally large, >540 kilobase (kbp) pairs in length (here referred to as “megaphage”). For context, as of June, 2016, only 93 phages with genomes >200 kbp (“jumbo phage”) were isolated, and none has a >500 kbp (13). The average length of complete phage genomes is 53,644 ± 45,677 bp, consistent with the average length of isolated dsDNA viruses of 44,296 ± 83,777 bp (14). We refer to the megaphage discovered in the Laksam Upazila cohort as Lak phage. Based on CRISPR spacer targeting, Lak phage are predicted to replicate in *Prevotella* species, bacteria that tend to be enriched in gut microbiomes of individuals who consume non-Western diets, as occurs frequently in the developing world (12). To determine whether these phage are common in other human and animal microbiomes, we assembled several DNA read datasets from samples containing abundant *Prevotella*. Faecal samples from the Hadza people of Tanzania (15),cholera patients from Bangladesh (4),Danish pigs, and members of two baboon social groups from Kenya (16) had evidence of Lak phage exposure or active infection, whereas samples from several unpublished datasets and Peruvian gut microbiomes (17) did not. Analysis of 15 Lak phage genomes that were manually curated to completion allowed us to investigate the megaphage diversity, distribution and genome dynamics. Overall, our results indicate that Lak phage are common and likely important components of gut microbiomes of humans and animals.

## RESULTS

### Megaphage identified in the gut microbiomes of Bangladeshi adults

DNA was extracted from either three or four faecal samples collected one day apart from 10 adults living in Eruani village, Laksam Upazila, Bangladesh, and sequenced (**Table S1**;see **Methods**). Taxonomic classifications and relative abundance information reveal the communities are mostly dominated by *Prevotella* species (**Figure 1**).

**Figure 1:**
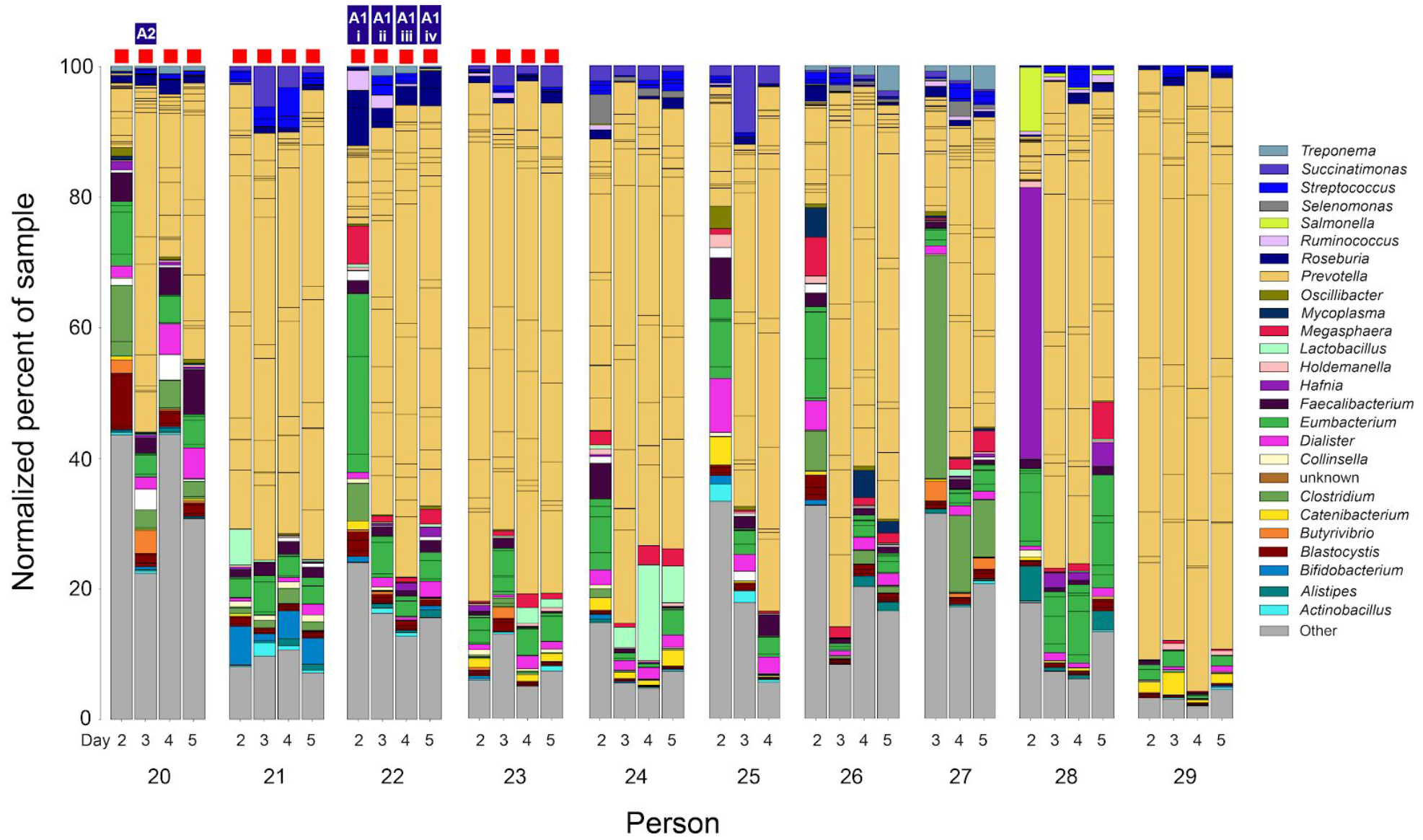
Community composition of gut microbiomes of ten subjects from Laksam Upazila, based on phylogenetic analysis of ribosomal protein S3 sequences. The increments in the stacked bar chart are colored by genus. Red boxes indicate samples from which megaphage sequences were partially assembled; blue boxes indicate samples from which a complete megaphage genome was recovered. Lines within each genus denote the relative abundance of genotypic variants.

Within assembled metagenomes from subjects 20 and 22 we identified large genome fragments that encode numerous hypothetical proteins, the majority of which have no similarity to any sequences in reference databases. These fragments encode phage genes, and thus were selected for manual assembly curation (**Methods**). Two bioinformatically verified, circularized phage genomes, A1 and A2, were >541 kbp in length, close to the maximum size ever reported for a phage (14). There was no evidence for integration of these sequences into bacterial genomes (**Supplementary Results**). Given their extraordinary size and to distinguish them from jumbo phage (>200 kbp genomes, (13)),we refer to these as “megaphage”. The A1 and A2 genomes are largely syntenic and share 91.3% average nucleotide identity (ANI). They encode ~35 putative tRNA genes (**Table 1**), many of which are concentrated in specific genomic regions. The megaphage were also detected in the other samples from Subjects 22 and 20. The A1 phage sequences from the three other Subject 22 samples were independently curated to completion to enable day-to-day and within-sample sequence variation analysis (**Table 1**). In all 10 regions where the four aligned A1 genomes differ, the distinct sequence occurred as a minor variant in all samples. Different site frequencies indicate that the variants are not shared by one or a few subdominant genotypes. Further, distinct patterns of change in relative abundances of population variants over time confirm that the polymorphic sites are not linked (**Figure S1**). Shifting relative abundances of subpopulations suggest ongoing replication over the four days.

**Table 1:** List of complete megaphage genomes. For additional information, see **Table S1**. ^+^Variants within incomplete genomes have tRNA introns not found in the C1 genome. Baboons are from two social groups, V = Viola’s and M = Mica’s (see (16)).

| Phage | Sample of origin | GC % | Length (bp) | # tRNAs | # tRNA introns | # Predicted ORFs (Code 15) |
| --- | --- | --- | --- | --- | --- | --- |
| A1-i | As cohort 22, #2 | 25.9 | 541643 | 33 | 3 | 581 |
| A1-ii | As cohort 22, #3 | 25.9 | 541664 | 33 | 3 | 581 |
| A1-iii | As cohort 22, #4 | 25.9 | 541664 | 33 | 3 | 584 |
| A1-iv | As cohort 22, #5 | 25.9 | 541664 | 33 | 3 | 581 |
| A2 | As cohort 20, #3 | 26.0 | 541299 | 34 | 4 | 581 |
| C1 | Cholera CH_A02_001D1 | 25.8 | 540217 | 32 | 2<sup>+</sup> | 591 |
| B1 | Baboon F22 (V) | 26.0 | 547991 | 30 | 1 | 591 |
| B2 | Baboon F3 (V) | 26.0 | 549839 | 31 | 1 | 594 |
| B3 | Baboon M09 (V) | 26.0 | 546746 | 30 | 1 | 590 |
| B4 | Baboon F30 (V) | 26.0 | 550552 | 31 | 1 | 594 |
| B5 | Baboon F18 (V) | 26.7 | 543529 | 31 | 1 | 583 |
| B6 | Baboon F16 (V) | 25.8 | 546689 | 30 | 1 | 588 |
| B7 | Baboon F11 (M) | 26.0 | 550702 | 31 | 1 | 599 |
| B8 | Baboon F4 (V) | 26.0 | 551627 | 31 | 1 | 600 |
| B9 | Baboon F01 (V) | 26.0 | 550053 | 30 | 1 | 593 |

### Prevotella are the host for the megaphage, based on CRISPR targeting

An important question is the identity of the host bacteria in which the megaphage replicate. Hosts can be identified based on CRISPR targeting because these immunity-relevant sequences are acquired during infection events (18). We extracted spacer sequences from all 2,485 CRISPR loci encoded on reconstructed genome fragments of >1000 bp in length. From 26 loci, 12 of which were in genomes of Prevotella species and 14 of which were on scaffolds too short to assign taxonomy to, 46 unique spacers targeted the A1 and A2 genomes. In many cases, spacers targeted both genomes, as expected given their high overall nucleotide similarity. Most of the Cas systems are Type I or Type 2 (with Cpf1, or Cas9) and spacers from both systems target the megaphage. Loci from these systems are known to undergo rapid expansion, and in some cases the *Prevotella* arrays contain up to 161 spacer repeat units. Most CRISPR loci show the typical pattern of shared spacers at one end and non-clonality associated with the currently diversifying CRISPR locus end. Spacers that target the megaphage occur throughout the loci (**Figures 2** **and** **S2**).

Only six (of 38) samples lacked *Prevotella* with CRISPR spacers that target the A1 and A2 phage, two from Subject 22 and all four samples from Subject 20. The lack of CRISPR spacer-based immunity may explain why the phage proliferated in those microbiomes. In fact, *Prevotella* species with spacers that target the megaphage occurred in samples from all subjects from which the phage was not reconstructed, consistent with the expectation that perfect spacer targets preclude infection at any appreciable level. Based on the prevalence of megaphage-targeting spacers, we conclude that these megaphage are common in the microbiomes of this human cohort.

Given that *Prevotella* species are the megaphage hosts, and many of the subjects have gut microbiomes dominated by *Prevotella*, we tested for megaphage in all the Laksam Upazila microbiome samples, and found evidence for them in samples from Subjects 21 and 23 (**Supplementary Information**). We attempted to isolate the megaphage using faecal material and *Prevotella copri* DSM 18205 (19)but isolation was unsuccessful (**Supplementary Information**).

**Figure 2:**
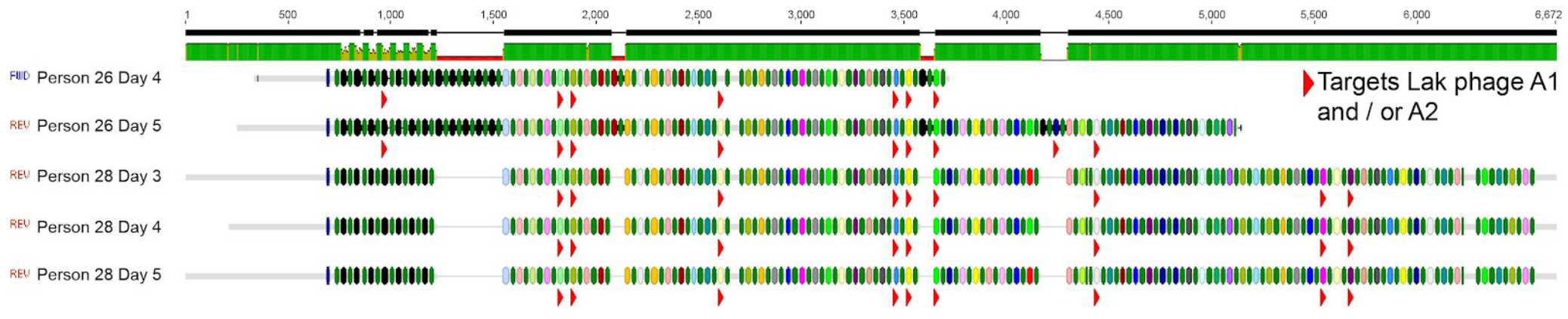
Diagram illustrating an alignment of the CRISPR arrays encoded on genome sequences of four *Prevotella* scaffolds containing repeat GGTTTAATCGTACCTTTATGGAATTGAAAT. Small green rods indicate repeats, colored rods indicate spacers. The same color indicates the same spacer sequence, except where the rods are black and indicate novel spacers in the datasets (novel spacers have likely been added to the diversifying ends of the loci). Gaps were inserted to align flanking regions and spacers with the same sequence and indicate regions where spacer-repeat array blocks apparently have been deleted. Red arrows appear under spacers that target the megaphage. For full version, see **Figure S2**.

### Megaphage occur in other gut microbiomes

Bacteria of the genus *Prevotella* are abundant in gut microbiomes of humans in the developing world. Thus, we wondered if related megaphage occur in other gut microbiomes that contain *Prevotella*. A search of NCBI’s non-redundant protein database for proteins related to those of the megaphage yielded no significant hits, so we selected individual metagenomic datasets from *Prevotella*-enriched samples for deeper analysis. In a prior study, faecal samples were collected from a cohort of Bangladeshi cholera patients and their families (42 male adults, 3 female adults, 2 male and 2 female children) who were hospitalized in Dhaka, Bangladesh in 2016, but the reads were not assembled (4). We conducted genome-resolved metagenomic analyses of these datasets. Many of the gut microbiomes were dominated by *Prevotella*, and, consistent with our prediction, they also contained phage related to the A1 and A2 megaphage. One 540,217 kbp genome, C1, was manually curated to completion. A dataset from a second Bangladeshi cohort comprising six cholera-impacted adults was sampled from the same hospital in 2011. Of those, three had significant (>5%) *Prevotella* populations and sample S75 had relatively abundant phage related to C1. Two other adults with low *Prevotella* (<0.1%), samples S71 and S72, had >100 reads map to the C1 genome (**Figure S3**).

Gut microbiome (faecal) samples from individuals from the Hadza tribe of Tanzania were sequenced in prior research (15). We mapped reads from 27 Hadza subjects to the megaphage genomes and found that three subjects (**Figure S3**)had these phage in sufficiently high abundance for genome assembly. The assemblies were highly fragmented but sequences shared ~90% identity to phage A1 (**Table S1**). A previously sequenced cohort of children living in India (20) also contained evidence of the megaphage, as two samples from this cohort had 500 bp reads that, in combination, covered more than 50 kbp of the A1 genome.

Previously published metagenomic shotgun sequencing datasets from faecal samples from 48 members of two social groups of Kenyan yellow baboons (*Papio cynocephalus*; one metagenome per individual) were assembled and investigated to identify putative megaphage sequences (16). Megaphage were detected in 43 of the 48 baboon gut microbiomes, and all samples contained multiple *Prevotella* strains or species (**Figure S4**). Sixteen high quality genome bins were identified from 16 distinct samples, nine of which were curated to completion (B1-B9). All genomes were >543 kbp in length, and one (B8) is the largest phage genome reported in this study (551,627 bp). All encode either 31 or 32 putative tRNAs (**Table 1**). The sets of tRNAs in the baboon megaphage genomes are more similar to each other than to the sets in the other megaphage genomes (**Table S2A**),although allowing for rearrangements (**Table S2B**), there are strong similarities in the tRNA complement across megaphage from all datasets.

To extend the search for the megaphage in other human and animal microbiomes we analyzed sequence data from *Prevotella*-containing samples from Danish pigs. Samples from pigs on the same farm were pooled and sequenced together (n=105 farms) (**Table S1**). Unfortunately, the genomes are highly fragmented, but based on sequence alignments, we identified a total of 18.7 Mbp of contiguous putative megaphage sequence with an alignment length 15.9 Mbp to the A1 genome. In one case, we identified a tentative partial genome bin that comprises 462 Kbp of sequence. At least 2 kbp of aligned megaphage sequence was detected in 104 of the 105 metagenomes. Over a 30 kbp alignment length, pig-derived sequences share an average of ~91.8% ANI with the A1 genome (**Figure S5**). Thus, we conclude that megaphage related to those present in humans and baboons also colonize pigs. We also found that reads from four of 27 cow rumen metagenomes (21) mapped with low coverage across the entire A1 phage genome (**Table S1**). However, analyses of 34 *Prevotella*-rich metagenomes from faecal samples from Tunapuco, a traditional agricultural community in the Andean highlands, did not detect megaphage (17).

To investigate whether samples that contain the megaphage share similar *Prevotella* species, we constructed a phylogenetic tree based on 16S rRNA gene sequences (**Figure S6**). Typically, the microbiomes contain a diversity of *Prevotella* strains, but overall, there was no clear link between cohort and *Prevotella* species or the presence of the megaphage.

### Megaphage use an alternative genetic code

A notable feature of the megaphage genomes was their low coding density (<70% for A1 and A2) when genes were predicted using the normal bacterial code (code 11). Investigation of the pattern of distribution of open reading frames and of protein sequences (some of which were clearly interrupted) indicated that the megaphage might be using an alternative genetic code. We tested a variety of ways in which a stop codon might have been repurposed to encode for an amino acid and determined that genetic code 15 is probably used by all of the megaphage reported here (TAG to glutamine, Q; **Figure S7**). Even after re-prediction with code 15, there are unusually large regions (up to ~1.5 kbp in length) that are not predicted to encode proteins or tRNAs.

To investigate megaphage activity *in situ* and to test for expression of the large intergenic regions we obtained metatranscriptomes from all samples from subjects 20 and 22. Thousands of transcript reads mapped to the A1 and A2 genomes (e.g., see **Figures S8**),indicating that at least a subset of the phage were replicating at the time of sample collection. The transcript mapping patterns support the use of an alternative genetic code and indicate that many intergenic regions are expressed. In fact, some intergenic regions are among the most highly expressed portions of the genomes (**Figures S8A-C**).

When the distribution of the repurposed TAG codon is plotted over the aligned A1 and A2 genomes it is apparent that this codon is not used in large regions (red bars in **Figure 3A**). However, it is used in some sections of the genome, including most that encode structural proteins (**Figures 3A** and **S8**). In samples from all days, we detected expression in regions encoding genes that do and do not use the TAG codon. Thus, if genes encoding structural proteins are expressed late, the phage in each sample are in a variety of stages of replication (**Figure S8D**).

Genomes in which a stop codon has been repurposed typically encode a suppressor tRNA (Sup
tRNA). Multiple types of Sup tRNAs were predicted (**Supplementary Information** and **Table S2**). The anticodon of one of these is CTA, which is necessary to repurpose the TAG stop codon. All complete megaphage genomes also encode Release Factor-2, which terminates translation by recognizing the TGA and TAA but not TAG stop codons. Thus, the megaphage have the cellular machinery necessary to successfully translate genes with in-frame re-coded TAG.

### Comparative genomics of the megaphage

One gene that is important during capsid assembly is the terminase (which was interrupted 18 times when proteins were predicted using genetic code 11). Based on phylogenetic analyses using terminase sequences (**Figure S9**),the megaphage are divergent and place generally within the Myoviridiae. Given that they are clearly a new clade and highly distinct in terms of their consistently very large genomes and use of alternative coding, we define them as the “Lak phage”. This name derives from the name of the region (i.e., Laksam Upazila, Bangladesh) from where the megaphage were first detected.

The A1, A2 and C1 genomes are syntenous, as are all baboon Lak (B-Lak) genomes, but six large rearrangements distinguish the B-Lak from the A-Lak and C1-Lak genomes (**Figure S10**). As expected based on their synteny, A1 and A2 are more similar to C1 than the B-Lak genomes (**Table S3**). Alignment of one ~70 kbp region from each genome with the A1 genome (**Figure 3B**)shows that insertions/deletions of sequence blocks within the central regions of genes (also see **Figure S11**), complete gene insertion/deletions, intergenic insertions/deletions and varying levels of nucleotide substitutions (sometimes varying greatly within a gene, **Figure S11**)distinguish the genomes.

Alignment of the nine B-Lak genomes revealed that the B-Lak genomes share ANI values of between 88.5% and 99.9% with each other (**Table S3**)and over the region in **Figure 3**,~61% to 65% ANI with A1. For comparison, A1 and A2 share ~95% ANI and A1 and C1 share ~88% ANI over this region (**Table S3**).

To compare the B-Lak genomes to each other, B1 was set as the reference sequence and B7 designated as the variant end member (**Figure S12**). A ~20 kbp region of this alignment is shown in **Figure 4A**. Bars underlining sequence blocks with the same color indicate regions with perfect sequence identity. Notably, identical sequence blocks varying in size from ~100 bp to tens of kbp occur in multiple phage genotypes and all genomes share some sequence blocks with the B7 genome. In some cases hundreds of shared single nucleotide polymorphisms (SNPs) distinguish sequences from the sequences of other genotypes. The strong signal of sequence block admixture clearly indicates reassortment. Few SNPs distinguish otherwise identical blocks. Some sequence blocks are unique to one of the B-Lak genomes, but overall the results indicate that the studied B-Lak phage have extensively exchanged genetic material. Presumably, recombination events require co-infection, i.e., the coexistence of these huge genomes inside a *Prevotella* cell at the same time.

**Figure 3:**
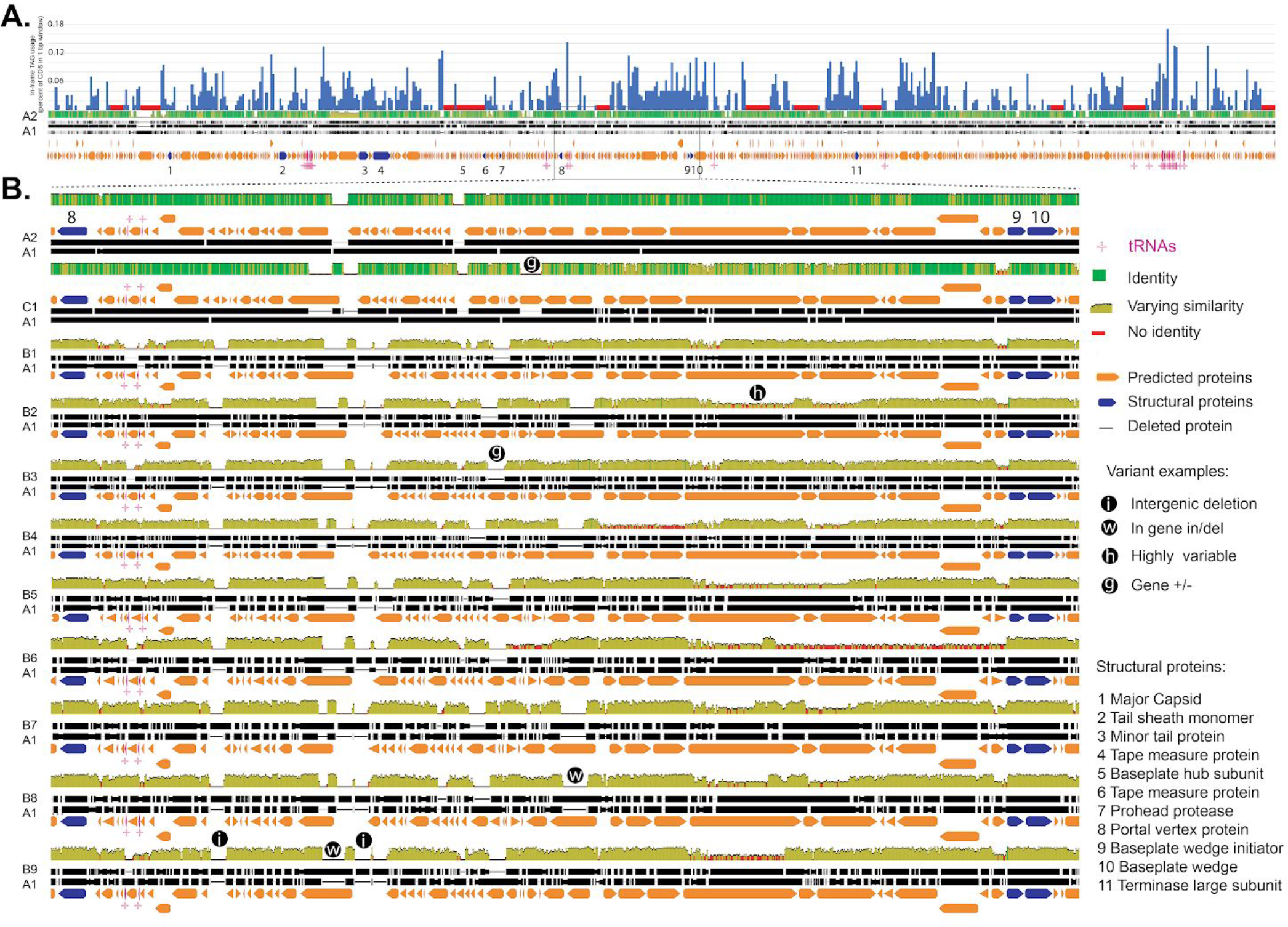
Genomic features and variation in Lak phage genomes. **A**. Frequency of use of the TAG stop codon, which has been repurposed to encode glutamine (top panel) overlying the alignment of the A1 and A2 Lak genomes. Red bars indicate regions >5kb with no in-frame TAG usage. Structural proteins (dark blue) are mostly encoded in regions with high in-frame TAG usage. The box indicates one largely syntenic region that is shown in detail in **B**.**B**. Alignments of each distinct Lak genome (excluding the A1 variants) against the A1 genome. Regions that could not be automatically aligned were manually aligned, and in some cases forcibly aligned, for clarity of presentation. A subset of this genomic region was used in the pig Lak genome fragment alignments (**Figure S5**).

**Figure 4:**
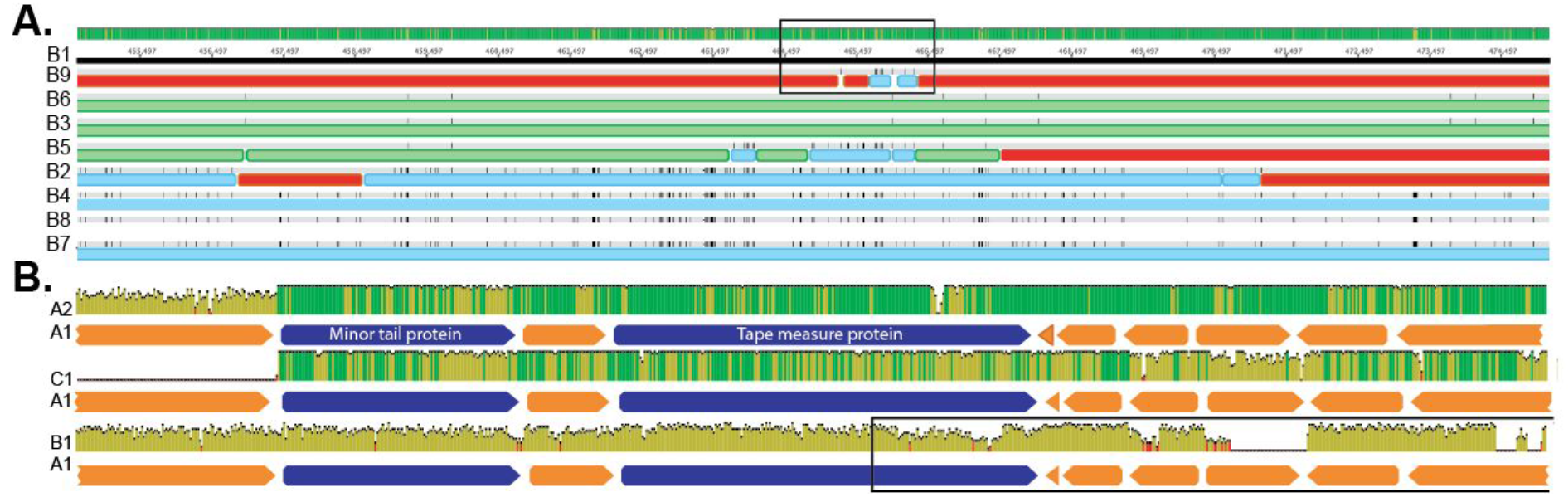
**A**. Sequence variation in a ~20 kbp region of the aligned B-Lak genomes with B1 as the reference sequence. The region includes part of a tape measure protein and subsequent predicted proteins. Colored bars designate sequence blocks, defined by single nucleotide polymorphisms. Note evidence of admixture of sequence blocks, indicative of extensive homologous recombination among phage sampled from individual baboons. For the full alignment of the nine complete Lak phage genomes, see **Figure S12**. The box indicates the region of the B9 population that was examined in detail (**Figure S14**). **B.** The locus shown in **A**,indicated by an open box, includes relatively conserved and more divergent regions in the genomes from the Laksam Upazila (A1 *vs.* A2), cholera-impacted (A1 *vs.* C1) and baboon (A1 *vs*. B1) cohorts. For the key to sequence identity plots, see **Figure 3**.

### Lak phage populations are near-clonal, but some contain sequences of other phage

To probe within-population heterogeneity and rule out the possibility that the patterns of scrambling of sequence motifs are artifacts due to the choice of a composite reference sequence, we analyzed sequence variation in reads mapped to each B-Lak genome (**Table S4**). In all cases, between 94% and 96.4% of the reads map to the genome with zero SNPs, providing confidence that the reported genomes are not chimeras of population variants. However, 0.01% to 0.8% of reads in each dataset match the sequences of other B-Lak genomes (**Table S4**). The majority of the reads with one or more SNP that cannot be linked to a known B-Lak phage sequence map throughout the genomes and probably reflect random sequence variation. However, a subset of the reads (especially from B4, B7 and B9) tile out variant sequences (probably recombinant genotypes, given the high degree of localization of these read mappings), likely derived from B-Lak genomes not reconstructed to date. In a few instances adjacent SNP groups within individual Illumina reads from the B9 dataset are linked and in other reads they are not (**Figure S13**),again indicative of reassortment of alleles *via* homologous recombination. As a result, there is no true meaning to a consensus genome sequence in this region.

From one region of the B9 population genomic dataset with an elevated frequency of reads carrying sequence variants (**Figure 4A** box) we extracted polymorphic reads and their paired reads (and a few read pairs without SNPs to illustrate support for the consensus sequence) and mapped them back to the B9 genome (**Figure S14**). Some paired read variants could be assigned to the B7 genome, extending backward from the small region where B9 and B7 are identical. Many variant reads could be assigned to B1, perhaps not surprising given the overall high similarity between the B1 and B9 genomes. Others could be assigned to B2 and/or B5. A subset of read pairs had SNP combinations that were distinct from those of known genomes, supporting the inference of recombination with as yet unreconstructed B-Lak genotypes.

The evidence of extensive homologous recombination motivated the question of whether phage relatedness (itself partially due to recombination) and phage admixture (sub-dominant genotypes within each population) could be predicted by baboon relatedness or social behavior (**Table S5**). Based on genetic (pedigree-based) estimates of kinship, grooming interactions (which have previously been linked to similarity in gut microbiomes in baboons, see (16)),and host spatial proximity, there is no strong indication that factors that could impact phage distribution across the baboon cohort have strongly influenced the current phage genomes or within-population variation (**Figure S15**).

### Lak phage encode tRNAs with highly conserved introns

An interesting feature of the Lak phage genomes is the presence of tRNAs with introns. To our knowledge, introns in phage tRNAs have not been reported previously. Several occur in A1, A2 and C1 tRNAs and one occurs in tRNA Tyr (GTA) in the baboon Lak phage genomes (**Table 1** and **Table S6**). The set of tRNA introns in the A1, A2 and C1 genomes only partially overlap. Intriguingly, however, where the same intron occurs in the same tRNA, its sequence is typically identical across cohorts (**Supplementary Information**).

We predicted the tRNA intron excision points using an implementation of tRNA scan designed for eukaryotes (as bacterial tRNA genes typically do not encode introns and so the program does not recognize them). Interestingly, the introns in putative tRNA Thr (TGT) are predicted to encode a possible tRNA (**Table S6**),the sequence of which is preserved perfectly in all but one case across cohorts. C1 and some other phage fragments from the cholera cohort lack this tRNA intron and alignment of sequences with and without this intron reveals that the excision occurs in a slightly different position than predicted (**Figure 5**).

The largest putatively circularized phage genome reported to date from any environment is 595,573 bp in length, with an unknown host (14). That genome is predicted to encode 75 tRNAs, 44 of which have unknown or mismatch isotypes. We predict that this genome has at least 4 tRNAs with introns. Thus, tRNAs with introns may be fairly common in large phage genomes.

**Figure 5:**
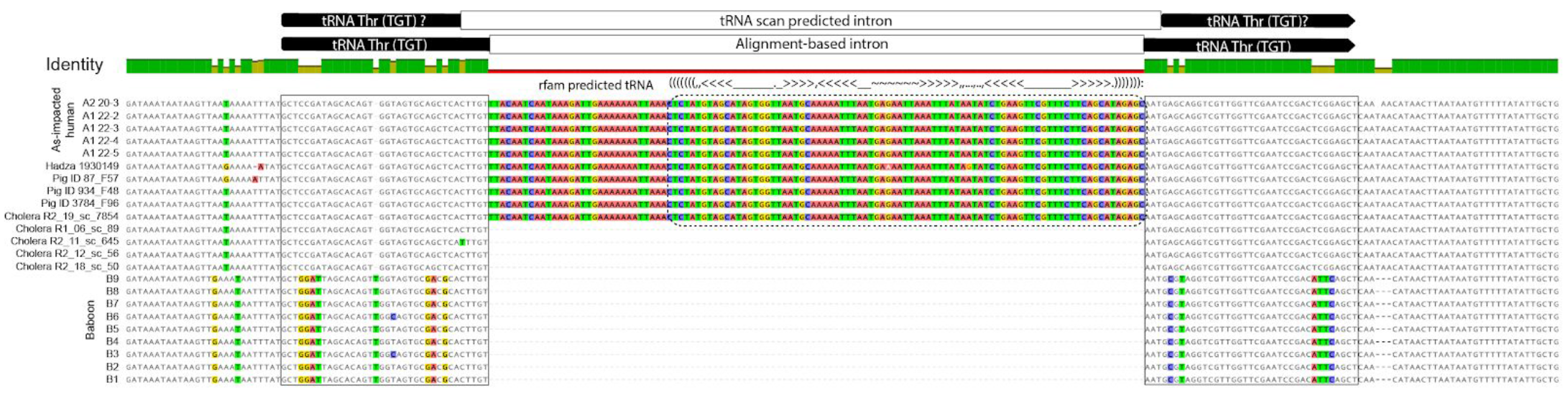
Alignment of sequences that contain a possible tRNA Thr (TGT). A subset of the tRNAs are predicted to contain an intron that itself is predicted to encode a possible tRNA (dashed box with superimposed secondary structure notations). Note that, with the exception of the Hadza sequence, all of the introns are identical and all but the baboon tRNAs are essentially identical.

### Lak phage metabolism and impacts on host population dynamics

The huge size of the Lak phage genomes motivated investigation of the encoded genes and their potential importance for *Prevotella* metabolism (**Table S7**). As expected, the vast majority of protein coding genes are hypothetical (84%). Repeat sequence analysis identified a few genes with similar amino acid sequences that were probably duplicated. The largest inventory of genes with tentatively recognizable functions are involved in nucleotide, DNA and RNA transformations, functions previously noted as prominent in large phage genomes (13). In general, the nucleic acid-related functions predicted for these genes are similar to those predicted for genes of Candidate Phyla Radiation bacteria, which are symbionts that seem to repurpose scavenged DNA for nucleic acid synthesis. The phage may augment host translational machinery using sigma factors, translation initiation factors (e.g., IF1), chain release factors as well as some genes that modify tRNAs (e.g., for adding a guanine nucleotide to the 5’ end of tRNA^His^,peptidyl-tRNA release, and CCA addition for tRNA maturation).

Intriguingly, the genomes encode one or two proteins predicted to hydrolyze a compound related to nucleoside 3′,5′-cyclic phosphate to nucleoside 5′-phosphate. Such enzymes might affect the cellular levels of the host’s cyclic second messengers. Potentially related, the genomes also encode a T4-encoded RNA ligase, which phage may use to repair tRNAs that are broken when a host defense mechanism produces 2′,3′-cyclic phosphate ends (22)

Among the enigmatic predicted proteins of potential significance from the perspective of host metabolism is an enzyme annotated as heme oxygenase [EC:1.14.99.3], which catalyzes the degradation of heme, an iron complex outer membrane receptor protein and a glycosyltransferase implicated in lipopolysaccharide biosynthesis. Despite the genome size however, we could recognize relatively few genes that might supplement host metabolism, except as it might pertain to phage replication.

Given that the exact host for a Lak phage cannot be established by CRISPR targeting if the *Prevotella* are susceptible to infection, it is difficult to confidently infer impacts of phage predation on microbial community structure. Thus, we compared the abundances of all (rather than specific) *Prevotella* species and Lak phage abundances over the four-sample Laksam Upazila time series. For both sample series, the phage are most abundant in the first sample and decline in abundance in the second sample with corresponding increases in abundances of the more abundant *Prevotella* genotypes (**Figure S16**).*Prevotella* species likely differ to some extent in their metabolic capacities and/or growth rates and/or nutrient preferences, and changes in relative abundances of species as well as their cumulative abundance could alter overall gut microbiome function. Based on the limited observations for two adults, shifts in *Prevotella* abundances due to phage predation may occur on the day to day time scale.

## DISCUSSION

### Lak phage are common yet overlooked members of gut microbiomes

Somewhat analogous to the recent finding of crAssphage that infect *Bacteroides*, bacteria commonly associated with humans in the developed world, here we report Lak phage that infect *Prevotella*, bacteria that are often abundant in the gut microbiomes of communities in the developing world where consumption of a high fiber, low fat, non-western diet is common (12). Interestingly, the Lak phage are also prominent in the microbiomes of a variety of animals. Notably, the 15 complete curated Lak phage genomes are >5X larger than the crAssphage genome (10) and only ~4.6X smaller than the genomes of their *Prevotella* hosts (**Supplementary Results**).

It is interesting to wonder why megaphage that infect *Prevotella* have been overlooked, and why megaphage of any kind have been so rarely described. The explanation may be two fold. First, as suggested previously, very large phage are difficult to isolate due to the physical restrictions on their mobility on plates used for plaque assays (23). Compounding this problem, *Prevotella* and Bacteriodetes in general are understudied as hosts (24). Interestingly, however, a huge *Prevotella* phage from the cow rumen was imaged almost two decades ago (25). The phage was reported to have an icosahedral head of ~120 nm in diameter and a long curved tail that was 30 nm wide and up to 800 nm long. To our knowledge, the largest isolated phage infects *Bacillus megaterium* and has a 497 kbp genome, a head diameter of 160 nm and tail length of 453 nm (13). It is not possible to establish a direct relationship between the head and tail length and genome size, but the imaged phage may be related to the Lak megaphage described here.

Assembly of metagenomic sequencing reads remains relatively uncommon and even when reads are assembled, most genomes are highly fragmented. In the current study, we used an assembly algorithm that includes a scaffolding step, and this brought to light a 540 kbp long genome sequence that could be circularized. Circularization provides confidence in the classification of the sequence as phage *vs.* prophage (which should transition into flanking genome). Phage structural proteins needed to confirm classification of the sequences were obscured by distant homology to known sequences and gene fragmentation in the initial annotation due to the use of the wrong genetic code. Recognition of alternative coding was complicated by the unavailability of code 15 in the NCBI genetic code listing and ORF finder application. This code has only once been inferred to be used by phage (26). The lack of obvious structural proteins could have precluded prior recognition of Lak phage sequences.

Based on a meta-analysis of public data, Paez-Espino et al. (2016) produced a dataset that included fragments classified as very large phage genomes. We note, however, that 11 of the sequences >200 kbp are artifactual composites of identical repeated sequences (**Figure S17**). This underscores the importance of curation and repeat analysis when reporting phage sequences. Despite this issue, we suspect that many more phage with very large genomes will be uncovered in future metagenomic analyses.

### Environmental distribution and dispersal of Lak phage

Have the Lak phage coevolved with their hosts (i.e., vertically) or has there been facile dispersal across animal habitat types? Some evolutionarily divergent hosts, humans *vs.* baboons, have less closely related Lak phage, and phage from geographically distinct human populations differ more than phage within a single human population. Although this might be evidence for co-evolution of phage and animal hosts, the presence of Lak phage genome fragments in pig microbiomes with high similarity to those in the human gut microbiomes serves as counter evidence (**Figure S5**). Further, we did not detect patterns of *Prevotella* speciation consistent with animal host specificity (**Figure S6**),so we suspect that Lak phage as well as their bacterial hosts may be actively dispersing across animal habitats.

A second and related question is the importance of the competing processes of genome divergence *vs.* the cohesive process of homologous recombination in Lak phage evolution. This could not be probed in cases where only one or two genomes were reconstructed from each environment. The baboon data are informative regarding this question, and clearly indicate extensive allele reassortment involving all of the analyzed baboon phage populations. This would have required extensive sharing of phage among the baboons. We infer that recombination events are recent, based on the low frequencies of SNPs that distinguish otherwise identical sequence blocks in different B-Lak genomes, and we suspect it is ongoing, given the presence of minor recombinant variants within some populations. Overall, the ability to detect admixture of genotypes suggests that distinct phage were brought into contact relatively recently, possibly following migration from another animal reservoir (potentially other baboon groups), and then began to recombine. A similar phenomenon was previously reported in bacterial genotypes (27). Consistent with recent introduction of the Lak phage is their prevalence in the baboon population and associated low level of CRISPR-based immunity.

If *Prevotella* and their huge phage migrate among animal and human microbiomes, they could carry with them genes that are relevant to human and animal health and the spread of disease. The concept of zoonotic viruses is well established, but there may be analogous phenomena involving phage. Phage can disseminate virulence factors between bacterial strains, including toxin-encoding genes responsible for many important diseases such as diphtheria, cholera, dysentery, botulism, food poisoning, scalded skin syndrome, necrotizing pneumonia or scarlet fever (9, 28) and propagate other genes of medical interest among animal reservoirs, such as those involved in antimicrobial resistance. The finding of related Lak phage in baboon, pig, cow and human populations suggests this possibility and the probability that it may occur is clearly increased where phage have huge genomes.

### Possible drivers of megaphage evolution

Are jumbo and megaphage the consequence of random local genome expansion events, or might there be stabilizing forces that converge on a specific genome length? Unlike most prior studies, we generated complete genomes for phage from multiple distinct cohorts. Thus, we could document a consistency of genome size of around 540 ‑ 550 kbp. We suspect a physical explanation for this narrow size range, possibly related to an advantage of particles in the colloid size range. For example, the gut habitat may favor particles of a certain dimension as size affects particle flocculation, attachment and retention in specific pore spaces. Clearly not all gut-associated phage are large so this could be, at best, a partial explanation (although we predict that there are many more large phage in animal guts than are currently recognized).

The existence of huge phage motivates the general question of the costs and benefits to the phage of large genomes and the feedbacks that drive the evolution of these genomes. Interestingly, the Lak phage genomes are in the size range of those of many putative bacterial and archaeal symbionts (e.g., candidate phyla radiation bacteria and DPANN archaea; (29)). Phage with large genomes probably transfer genes to their hosts at a faster rate than phage with small genomes, which might improve the fitness of the host population, and thus its phage. However, this effect is probably small and may occur against the background of sequential acquisition of genes (and gene segments, see **Figure S11**)that confer advantage to the phage. Lak genomes encode many tRNAs, which could affect their lifestyle in many ways (see above). We note, however, that the span of genome used to encode tRNAs is small, motivating consideration of the possibility that the many hypothetical proteins predicted in the hundreds of kbp of Lak phage genome sequences somehow ensure successful phage replication in the face of host defense mechanisms. These genes could also be important for increasing the host range.

Evolution of large phage genomes, and thus few expensive particles per replication cycle, could be an ecological strategy analogous to K *vs.* R selection. Phage would normally be viewed as R strategists, leveraging the advantage of many offspring to ensure high probability that a particle will find a host in which it can replicate before loss of viability. For large phage, the countering trade off of a shift towards K selection could be improved survival as the result of the large capsid size, potentially due to increased stability of larger capsids, for example due to their smaller radius of curvature (30). Clearly, many factors could come into play, and direct experiments involving isolated phage and their hosts will be required to understand the intriguing phenomenon of megaphage in human and other animal gut microbiomes.

## CONCLUSION

A new type of megaphage are overlooked members of human and animal gut microbiomes. Their existence substantially increases the representation of phage whose genetic repertoires blur the boundaries that separate bacteria, bacterial symbionts and parasites / mobile elements. Their genomes hint at fascinating biology and as yet unexplored complexity in the dynamics of gut microbiomes.

## Acknowledgements

We thank Prof. Todd Lowe for helpful discussion related to tRNAs and introns and Dr. David Paez-Espino and colleagues at the Joint Genome Institute for generating sequences in a meta-analysis related to IMG-VR that enabled identification of the baboon cohort as a potential source of Lak phage sequences. We thank Dr. Shufei Lei and Katherine Lane for ggKbase support and Dr. Luis Barreiro for his contributions to the baboon study. Funding was provided by the National Institutes of Health (RAI092531A) and Sloan Foundation (G 2012-10-05) and to JFB, Chan Zuckerberg Institute Biohub funding to JFB, PT and KS, and NSF grant IOS 1053461 to EAA. We acknowledge (15),(20),(21) and (17) whose published research generated, respectively, the Hadza, Indian children, cow rumen, and Peruvian gut read datasets used in this study.

Faecal samples were collected from patients in the clinical Phase I/II SEASP trial in Bangladesh that was jointly led by Prof. Graham George and Prof. Ingrid Pickering (University of Saskatchewan), with the assistance of the SEASP team https://clinicaltrials.gov/ct2/show/NCT02377635, and funded by the Canadian Federal Government, through Grand Challenges Canada, Stars in Global Health and by the Global Institute for Water Security. We thank Dr. Olena Ponomarenko and Sanjit Shaha for organising transport of the faecal samples.

## Author contributions

The initial study was designed by JMS and JFB and refocused by AD, JFB and JMS. JMS isolated the nucleic acids from the Laksam Upazila Bangladesh cohort and provided the DNA sequencing. KS provided the DNA sequencing for the second cholera-impacted cohort. JFB and AD curated the genomes. AD constructed the phylogenetic trees and analyzed their codon use, with input from BCT and KA. AD and JFB conducted the comparative genomic analyses, with input from MO. AD and JFB analyzed the predicted protein sequences. JMS attempted the Lak phage isolations. PM and FA provided the pig metagenomic data, which was analyzed by PM and JFB. JT, EA, LB and RB generated the previously published baboon reads and provided input to the metadata analysis. PJT generated the previously published cholera-impacted cohort read dataset. JFB and AD wrote the manuscript, with input from JMS, MO and PM. All authors read and approved the manuscript.

## METHODS

### Samples, DNA and RNA extractions, sequencing, read archive analysis

Faecal samples were obtained from 10 Bangladeshi men (aged between 27-52) living in the Eruani village, Laksam Upazila, Bangladesh (samples were taken in April, 2016). All subjects displayed signs of arsenicosis and were consuming arsenic-contaminated drinking water. Samples were collected on four consecutive days (labelled days 2-5) and stored at −20°C until they were shipped to the Santini laboratory, UK, on dry ice. Samples were stored at −80°C until nucleic acid extractions were performed. DNA was isolated using the PowerFaecal DNA isolation kit (MoBio) according to the manufacturer’s instructions and stored at −20°C. DNA samples were sent to RTLGENOMICS (Texas, USA) on dry ice and prepared for sequencing using the Kapa HyperPlus Kit (Kapa Biosystems) following the manufacturer’s protocol, except that the DNA was fragmented physically using the Diagenode Bioruptor, instead of enzymatically. The resulting individual libraries were run on a Fragment Analyzer (Advanced Analytical) to assess the size distributions of the libraries, quantified using a Qubit 2.0 fluorometer (Life Technologies), and also quantified using the Kapa Library Quantification Kit (Kapa Biosystems). Individual libraries were then pooled equimolar into their respective lanes and loaded onto an Illumina HiSeq 2500 (Illumina, Inc.) 2 × 125 flow cell and sequenced. RNA was extracted from all four samples from subjects 20 and 22 with the Powermicrobiome RNA isolation kit (MoBio) according to the manufacturer’s instructuctions and stored at −80°C until they were shipped to the QB3 center at UC Berkeley on dry ice for sequencing.

Read datasets from previously published studies were selected based on the sampled environment and information about *Prevotella* content and downloaded from NCBI’s single read archive (SRA). Reads were mapped to the Lak phage genomes initially assembled from the Laksam Upazila cohort to determine whether or not Lak phage was present in the sample. Selected read sets were trimmed using sickle (https://github.com/najoshi/sickle) and each dataset assembled separately using IDBA-UD (31) with default parameters.

### Binning of draft genomes, genome curation and annotation

Bins were constructed from scaffolds of >1 kbp in length based on the combination of genome GC content, coverage and a phylogenetic profile. The phylogenetic profile was established based on gene-by-gene comparison to a reference genome dataset (31). We identified all sequences that encoded ribosomal protein S3, a gene that occurs in a relatively conserved block of genes that encode ribosomal proteins, and used these sequences to profile the overall community composition (taxonomy and abundance). Putative phage scaffolds were identified based on the high fraction of proteins with no related sequence in the database or similarity to phage proteins, as well as the presence of genes encoding structural proteins. Very large genome fragments were selected for curation. In cases where these were substantially shorter than the final genome length, candidate fragment collections identified based on consistency of GC content, coverage and phylogenetic profile were subject to curation. Coverage values were determined by read mapping using Bowtie2 (32) with default parameters for paired-reads.

The first genome curation step involved identification of local assembly errors and either correction of the errors or gap insertion using ra2.py (33). Curation of each genome was conducted independently and involved correction of local scaffolding errors and gaps, contig extension to enable joins and circularization, with manual resolution of regions of confusion. Reads from the sample were mapped to the scaffold assembled from that sample and unplaced paired reads used to extend ends and fill gaps. The curation was conducted in Geneious (34). Regions of confusion were identified based on much longer than expected placement of paired reads or backwards mapping of paired reads. Reads were manually relocated and reoriented. The final curated sequences were visualized throughout to confirm complete and accurate coverage of each genome. Final genomes were checked to confirm the absence of large repeated sequences that could have confounded the assembly. The start position was chosen in a random region so as to not interrupt a gene. Later reconstructed genomes were adjusted so that the start positions corresponded to those of earlier assembled genomes.

Genes were predicted on scaffolds >1 kbp using Prodigal (35),initially using genetic code 11. Subsequently, the Lak phage genes were re-predicted using code 15. Initial functional predictions were established based on similarity searches conducted using BLAST against UniProtKB and UniRef100 (36),and the Kyoto Encyclopedia of Genes and Genomes (KEGG) (37) and uploaded to ggKbase (https://ggkbase.berkeley.edu/project_groups/megaphage). In addition, genes were annotated by scanning *via* hmmsearch (38) with a collection of KEGG Hidden Markov Models (HMMs) representative of KEGG orthologous groups. The majority of phage structural genes were not identified in our initial functional predictions. Thus, we searched for proteins with unknown functions against the NCBI non-redundant protein database using BLAST-psi to identify remote homologs. These sequences were then aligned and searched against the Pfam_A_v31.0 and TIGRFAMs_v15.0 databases using HHPred to assign functions (39). The amino acid sequences of terminases were so divergent from terminases of previously analyzed phage that they could not be identified using standard functional prediction methods. We found the terminase gene by searching all predicted phage proteins against a protein database of terminase large subunits, taken from phage spanning several different families and including all identified terminase large subunit (TerL) proteins from phage with genomes of >200 kbp in length.

The tRNA genes were predicted using tRNA scan with bacterial settings (40). The tRNAs that were larger than expected and could not be assigned a classification were evaluated in terms of potential introns. Genes were re-predicted using the eukaryote settings to identify the tRNA type, anticodon and intron sequence. Intron excision points were re-evaluated based on alignments of genes with and without inserted sequences. Intron sequences were tested for possible classification using rfam (41).

Codon usage was calculated in consecutive 1 kb windows, and was reported as the percent of a specific codon out of all codons in the coding region of that window. Calculations were done using a custom script, cu.py (https://github.com/banfieldlab/cu_scripts).

### CRISPR targeting analyses

CRISPR arrays were predicted on all scaffolds >1kb in the Laksam Upazila cohort and the baboon cohort using a command line version of the program CRISPR Detect (42) with parameter-array_quality_score_cutoff=3. Only arrays with a score above the cutoff of 3 were considered. Spacers and repeat regions were extracted from the output files, and all spacers and repeats were searched against the Lak phage genomes A1 and A2 using BLASTN with parameter-task=short. No repeat regions had a hit to A1 or A2, so all spacers with a hit containing ≤ 1 mismatches and a length >24bp were considered to target Lak phage. The taxonomy of the scaffolds containing the CRISPR arrays with spacers targeting a Lak genome were determined by assigning taxonomy to all genes on the scaffold based on a similarity search using BLAST against the UniProtKB database. Scaffold taxonomy was assigned according to the highest taxonomic level shared by at least 50% of the genes on the scaffold. CRISPR arrays containing the repeat GGTTTAATCGTACCTTTATGGAATTGAAAT were chosen for reconstruction based on their high number of spacers targeting Lak. Arrays were manually aligned and spacers colored using Geneious (34).

### Phylogenetic and community compositional analyses

Community composition (**Figure 1** and **Figure S4**)was determined by read mapping to the conserved ribosomal protein S3 gene (*rps3*). The *rps3* genes were identified on all scaffolds >1kb in the Laksam Upazila Bangladeshi cohort and the Baboon cohort, and classified to the species level based on usearch clustering (43) with annotated proteins in the uniprot database. All *rps3* genes were then clustered at 90% identity using usearch, and a representative sequence from each cluster was chosen. Reads from each sample (all 10 people, 3 or 4 samples from consecutive days per person for the Bangladesh cohort and all 48 baboons) were mapped to these representative sequences, and the percent coverage of each *rps3* gene was determined. Percent coverage was then normalized by the sequencing depth of each sample to determine percent project. Any genus that was present in <10% cumulative abundance across all samples was grouped into the “other” category.

The *Prevotella* phylogenetic tree was constructed using 16S rRNA gene sequences. First, the Greengenes database (44) of complete 16S rRNA gene sequences was augmented with all 16S rRNA gene sequences from *Prevotella* reference sequences on NCBI that were independently classified as *Prevotella* (15 were assigned to a genus other than *Prevotella* and discarded). This augmented database was then used to classify 16S rRNA gene sequences from all samples in each study where the megaphage was found, including samples in publically available studies, using the assign_taxonomy.py script from qiime1 and default parameters (45). Sequences classified as *Prevotella* were aligned with all known reference *Prevotella* 16S rRNA gene sequences and an *Escherichia coli* 16S rRNA gene outgroup (NCBI ref. J01859.1) using MUSCLE (46). A tree was generated using RAxML-HPC2 on XSEDE (47) on the CIPRES web server (48) using parameters raxmlHPC-HYBRID -T 4 -n result -s infile.txt -m GTRGAMMA -p 12345 -k -f a -N 100 -x 12345 --asc-corr lewis. The tree was edited and annotated using iTOL (49).

The terminase large subunit gene (*terL*) was detected in all 15 Lak phage genomes using a BLASTP search against a database representing TerL proteins from a wide variety of phage and all known jumbo phage genomes in the NCBI database. The Lak TerL proteins were aligned along with the reference TerL proteins from jumbo phage using MUSCLE (46),and the tree was generated using RAxML-HPC2 on XSEDE (47) on the CIPRES web server (48) using parameters raxmlHPC-HYBRID -T 4 -n result -s infile.txt -p 12345 -m PROTGAMMADAYHOFF -f a -N 100 -x 12345 --asc-corr lewis. The tree was rooted with the terminase small subunit of *Agrobacterium* phage 7-7-1 (NC_019519.1). The tree was edited and annotated using iTol (49)

### Comparative genomics

Genome sequences were aligned using the progressiveMauve algorithm using default parameters (50). In certain regions the sequences were offset as the algorithm failed to align them. In some cases this could be corrected based on visual inspection. In other cases the sequences were superimposed to constrain the overall alignment length (very low similarity scores were then displayed).

### Pig genome fragment analysis

To search 105 metagenomes that were constructed from collections of faecal samples from Danish pig farms we aligned all the assembled pig metagenomic scaffolds of >1 kbp in length against the Lak phage A1 reference genome using NUCmer (Mummer 4.0.0beta). Filtering was done, so that alignments of at least 2 kbp and 70% nucleotide identity were kept. The lengths of scaffolds meeting these criteria were summed to estimate the total genome sequence attributable to Lak phage, and the total alignment length calculated. A metagenome was considered to contain Lak phage as long as at least one scaffold with an alignment length >2 kbp was identified.

***Data availability*:** The 15 Lak phage genomes have been deposited at NCBI under accession # (TBA). The genomes can also be downloaded from https://ggkbase.berkeley.edu/project_groups/megaphage. Read archive and other accession information is provided in **Table S1**.

### Statement of ethics

The human faecal samples obtained were part of a clinical Phase I/II study in rural Bangladesh entitled “Selenium and arsenic pharmacodynamics” (SEASP) run by Graham George (University of Saskatchewan) and funded by the Canadian Federal Government, through a program entitled Grand Challenges Canada, Stars in Global Health, with additional funds from the Global Institute for Water Security at the University of Saskatchewan. The SEASP trial was approved by the University of Saskatchewan Research Ethics Board (14-284) and the Bangladesh Medical Research Council (940,BMRC/NREC/2010-2013/291). Additional ethics approval was also obtained by UCL (7591/001).

## SUPPLEMENTARY RESULTS

### No evidence for megaphage as prophage

No reads or read pairs connected the megaphage scaffolds to bacterial scaffolds, and no phage fragments were found in bacterial scaffolds in the newly generated dataset. We checked all NCBI *Prevotella* genomes for fragments of the phage using MUMmer (51). The only finding was a ~60 bp region identical in several *Prevotella* genomes and A1. This small region occurred at the end of a transposon *Prevotella* and in an intergenic region in the phage genome. Thus, to date there is no evidence that the megaphage integrate into *Prevotella* genomes. For the 44 *Prevotella* genomes reconstructed from the baboon datasets, the median size was 2.38 Mbp (based on the cumulative length of scaffolds assigned to *Prevotella* genome bins), with a median of 50/51 expected single copy genes and 0.5 duplicated single copy genes per genome. For examples, see: https://ggkbase.berkeley.edu/project_groups/megaphage.

### Detection of phage in other subjects of the Laksam Upazila cohort

In Subject 21, co-assembly of reads mapping to phage A1 from all four samples resulted in 195,292 bp of contiguous sequences, the largest of which were around 900 bp in length and comprised of perfectly mapping reads. There was also read mapping across the entire A1 genome from samples from Subject 23, but the coverage was too low for assembly. All other samples had fewer than 250 reads mapping non-specifically to the A1 genome.

### Phage isolation attempts

We attempted to isolate the phage using faecal samples from subjects 20 and 22 following a similar method described previously (10).*Prevotella copri* (DSM 18206^T^was used as the host and grown in liquid and on solid media as described previously (19). In brief, faecal samples (about 1 g) were suspended in 1 ml buffer, vortexed, centrifuged to remove particulate material and filtered through a 0.45 μm filter. Confirmation that the Lak phage was present in the extract was done by PCR using Lak-specific primers. Mid-exponential phase *P. copri* cultures were used to inoculate plates (19) which were then seeded with the phage extract, incubated and then visualised for cell lysis. As was the case in the prior crAssphage study, isolation was not achieved, which may be due to a number of reasons including that *P. copri* is not a Lak phage host.

### Sup tRNA analysis

Questionable tRNA are recognized when the isotype predicted based on the overall tRNA sequence alignment does not match that predicted by the anticodon. In the case of Sup tRNA a mutation has occurred in the anticodon so that it now matches a stop codon. The tRNA scan program predicted glutamine as the third most likely amino acid linked to the Sup CTA tRNA, but this prediction is unreliable because the phage sequences are far distant from those used to build the models (Todd Lowe, pers. comm). The relevance of the other Sup tRNAs is uncertain.

### Detection of Lak phage in other Baboon metagenomes

Partial genomes were identified in the following Baboon metagenomic datasets:

1747022 (M05) ~540 kbp in six scaffolds, largest is 230 kbp.
1747029 (M06) ~537 kbp in three scaffolds, largest is 432 kbp.
1747031 (M08) ~546 kbp in two scaffolds, larest is 361 kbp
1747033 (F07) ~545 kbp in five fragments, largest is 281 kbp.
1747038 (F12) ~546 kbp in two scaffolds, largest is 361 kbp.
1747039 (M10) ~ 541kbp in four scaffolds,largest is 151 kbp.
1747060 (F28) ~ 546 kbp in two scaffolds, largest is 361 kbp.

Bins are available: https://ggkbase.berkeley.edu/project_groups/megaphage

### B-Lak population variation analysis

As documented in **Table S4**,the B1, B2, B3 and B5 populations are relatively clonal, and ~96.4% of reads that mapped to the genome using standard parameters can be mapped with zero SNPs. For these phage, 0.01% ‑ 0.02% of the reads map perfectly to another B-Lak genome. The B6 and B8 populations are only slightly less clonal and show similar fractions of reads with perfect matches to other B-Lak genomes (0.01 and 0.4% of all reads). Even for the other populations, B4, B7 and B9, between 94% and 94.7% of reads match perfectly to the reconstructed sequence, again providing confidence that the reported genomes are not chimeras of population variants.

Populations datasets for B4, B6 and B9 provide the best evidence for the presence of sub-dominant sequence types characteristic of another B-Lak population (probably as recombinant genotypes, given the high degree of localization of the read mappings). More than ~0.1% of reads from some population datasets map to other genomes (B4 reads to the B2, B8 and B9 genomes; B6 reads to the B3 and B9 genomes and 9 reads to five other genomes). The B9 genome is notable in that it was reconstructed from five genome fragments that comprised ~99% of the final bin length. Fragmentation was at least partially due to a few local regions of within-population sequence variation.

### tRNA intron similarity across cohorts

Introns were identified within some pseudo tRNAs that are substantially larger than expected. An identical tRNA Phe (GAA) intron is found in A1, A2, the cholera cohort and some pigs; identical introns occur in a tRNA Gly (TCC) in the A2, C1 and some fragments from pigs and identical tRNA Thr (TGT) introns occur in A1, A2, C1 and some genome fragments from the cholera and pig cohorts. Finally, we identified identical introns in Tyr (GTA) from A1, A2 and C1 and related but longer introns in the B-Lak phage genomes (**see Table S6**).

### Phage metabolic potential

The largest fraction of genes with confident functional predictions are involved in reactions involving nucleotides (See **Table S7**). Some genes encode enzymes that interconvert various deoxyribonucleoside and ribonucleosides, phosphorylate/dephosphorylate nucleotides and participate in other steps in DNA and RNA metabolism (e.g., ribose-5P to PRPP). In addition, there are helicases, topoisomerases, polymerase subunits, primases, exonucleases, genes involved in restriction, base excision and repair and recombination (potentially of relevance given the extensive evidence for homologous recombination).

The genomes encode several genes involved in one carbon metabolism and genes involved in interconversion of nicotinate and nicotinamide, possibly with an impact on host energy metabolism. Also predicted is pnuC, a nicotinamide mononucleotide transporter. Other genes involved in transport include a co-located three subunit NitT/TauT (sulfonate/nitrate/taurine) system. The genomes also typically encode multiple molecular chaperones.

### Infection cycle

Early in infection, the presence of release factor 1 (RF1, which would recognize UAG as the termination codon)would prevent production of structural proteins (as they would be fragmented). Given the very large phage genome size, its translation would deplete the cellular pool of RF factors. With Sup tRNA, and once RF2 becomes abundant, the phage genome can be correctly transcribed. If code 15 becomes important only relatively late in the phage replication process (26),transcriptomic data could have been informative regarding the stage of Lak phage replication if the phage replication cycle was synchronized **Figure S8**). We did not detect a clear signal for shifting replication phase, suggesting ongoing and uncoordinated replication events. Consistent with a more complex pattern of phage replication were small shifts in phage population structure over the four day period (**Figure S1**).

Phage-encoded tRNAs may correspond to codons that are abundant in certain phage genes and thus increase their translation efficiency (52). They may also decrease the probability of mismatches between host and phage codon preference and thus increase host range (13). The tRNAs and pseudo-tRNAs may have other purposes, however. For example, they may confuse the translational apparatus through loading of the wrong amino acid or stall translation due to inability to load amino acids.

## Supplementary Figure Legends

**Supplementary Figure 1.**
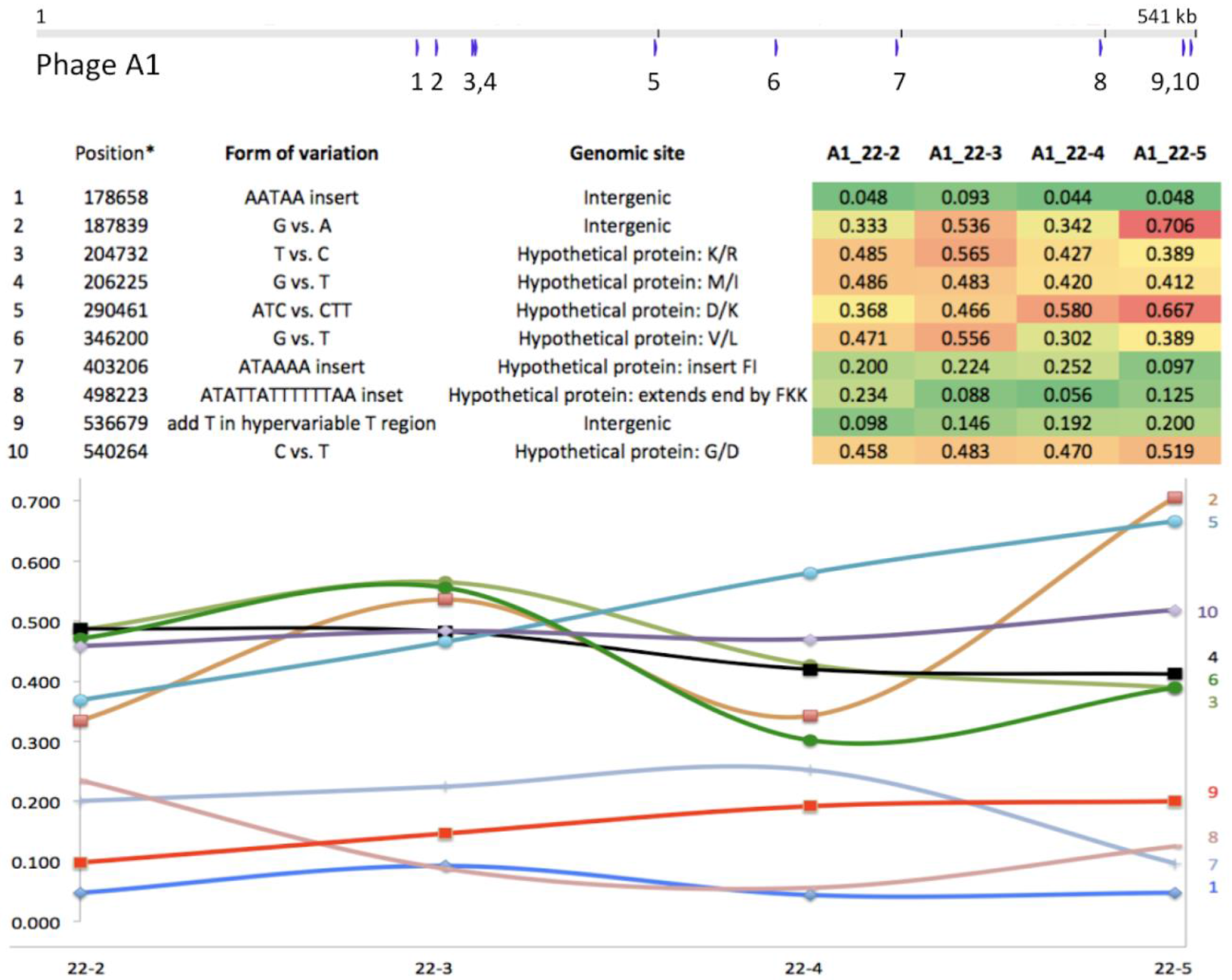
Analysis of the frequency of variants at ten variable sites (numbered blue tick marks) in the genomes of the Lak phage A1 in samples collected over four sequential days (22-2, 3, 4, 5). The table lists the variant site types, consequences of variation and site frequencies. The graph shows how the relative abundance of each variant changed over the sample series. Differences in the frequencies of variants at each site in a single dataset in combination with divergent patterns of variant frequencies over the sample series suggest that few if any of the variants are linked.

**Supplementary Figure 2.**
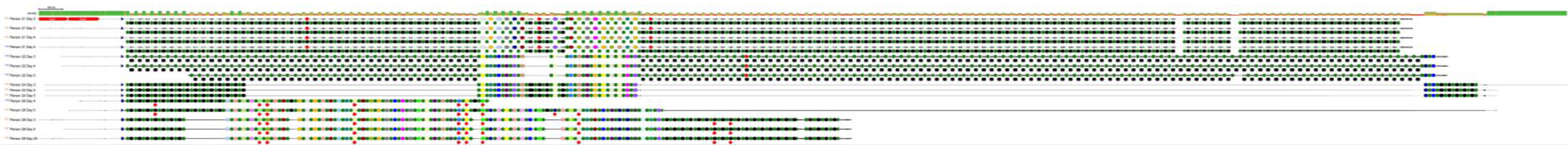
Full reconstruction of CRISPR arrays targeting megaphage from subjects 21, 22, 24, 26, and 28. Note that the *Prevotella* arrays from sample 22, where the phage is abundant and was first identified, has only one spacer targeting the megaphage. For details, see **Figure 2**.

**Supplementary Figure 3.**
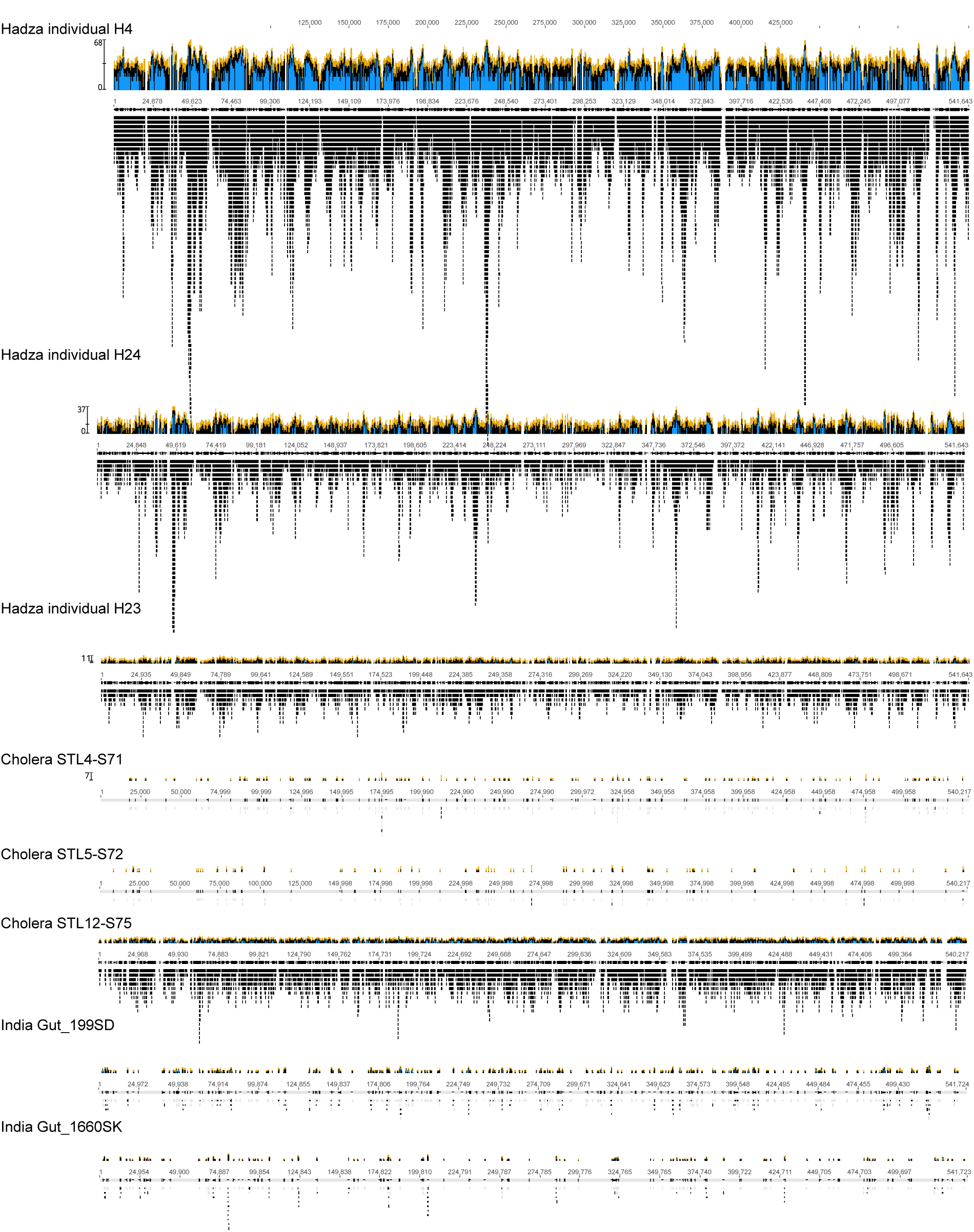
Overview of reads mapped to the Lak phage A1 genome from various cohorts. The coverage scale is approximately even throughout the presentation. For data accession numbers, see **Table S1**. Data sources are as follows: Hadza, (15); Cholera, (4); India, (20).

**Supplementary Figure 4.**
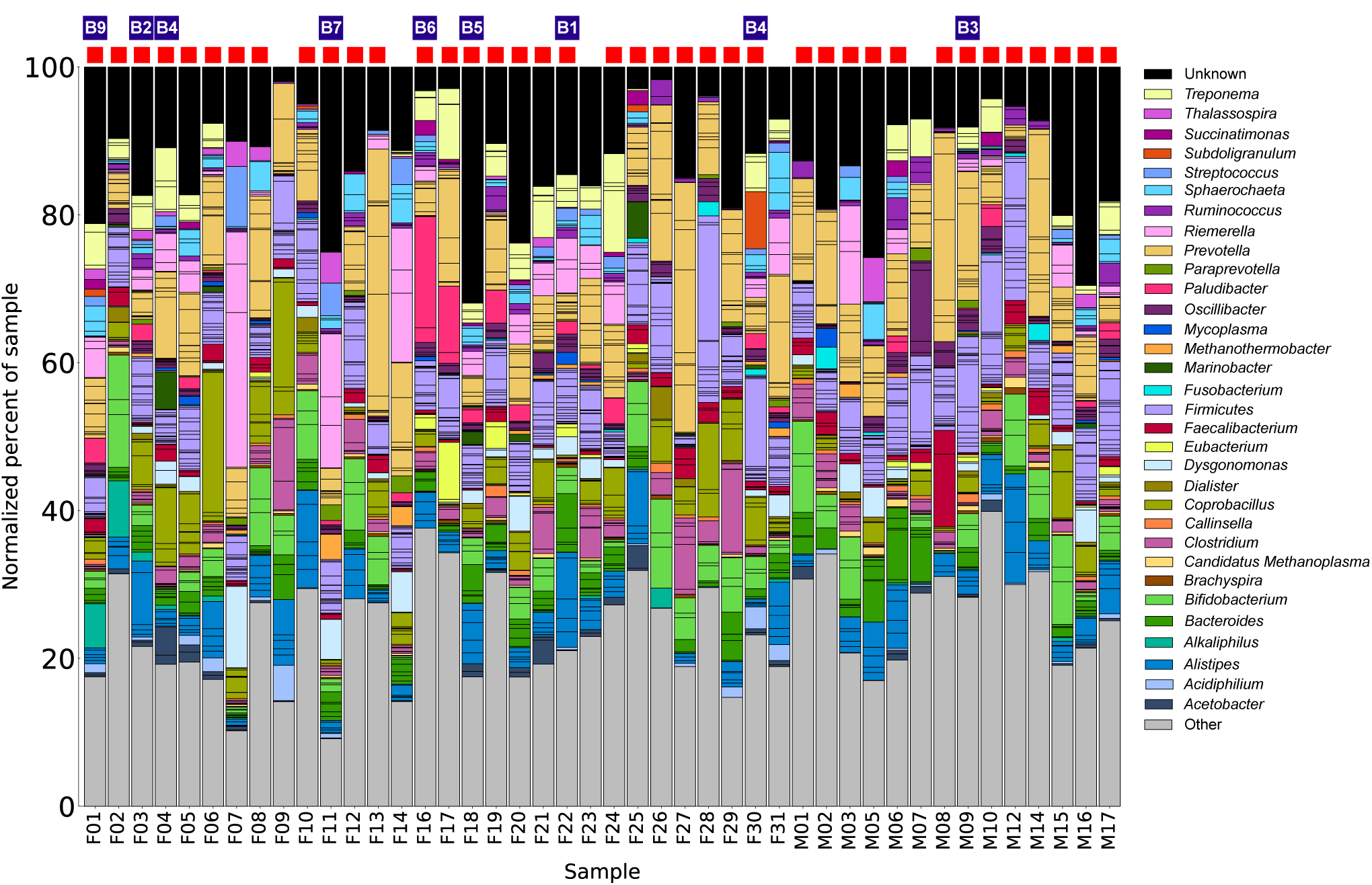
Overview of the community composition of baboon faecal samples. Red squares indicate samples from which Lak phage were partially assembled, blue squares indicate samples from which complete Lak genomes were reconstructed. Baboon sequencing data first reported by (16).

**Supplementary Figure 5.**
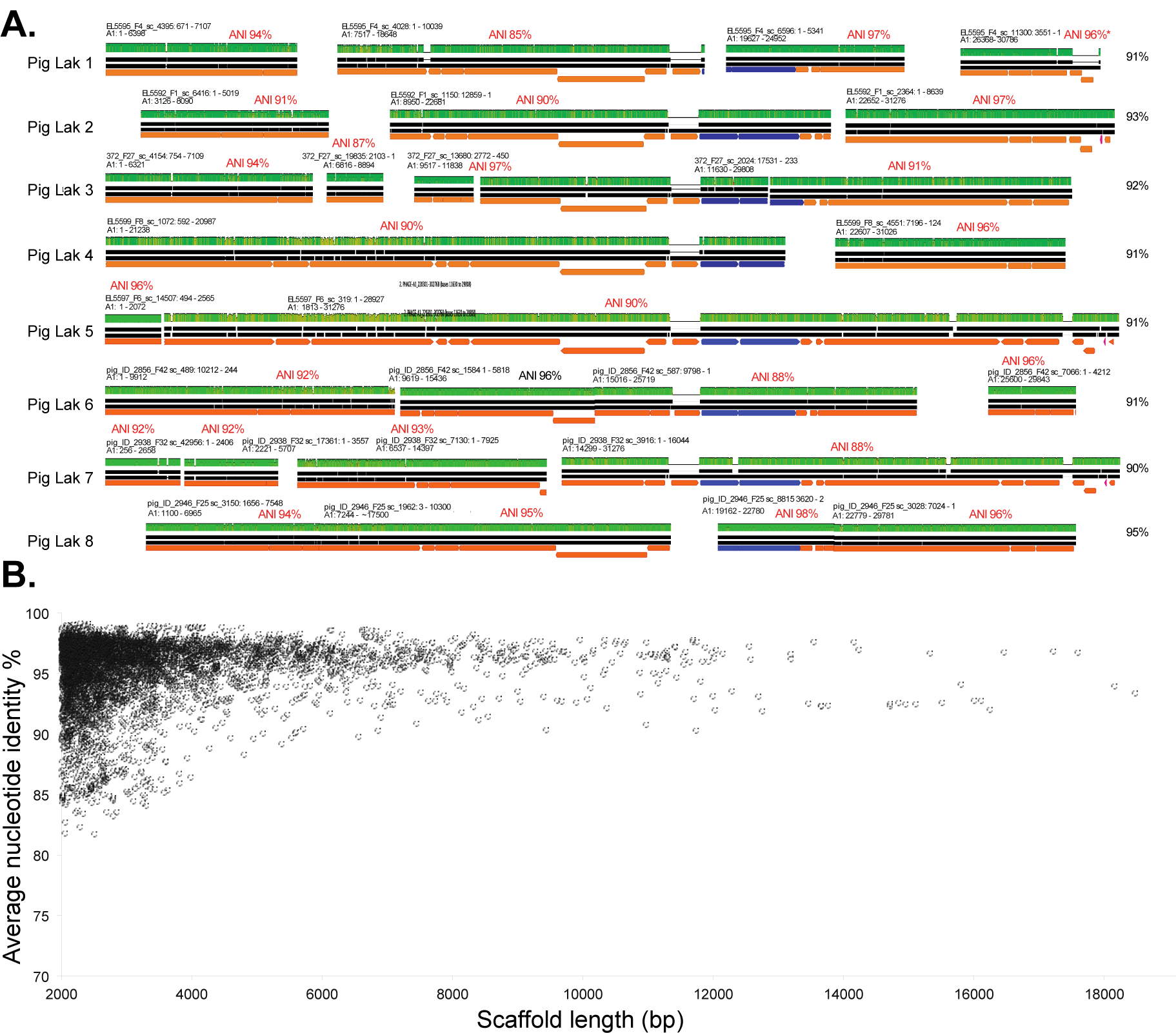
**A.** Fragments assembled from pig farm metagenome datasets were identified as deriving from Lak phage based on their sequence identity to the A1 Lak phage genome. The figure shows eight examples. Given that most fragments were <10 kbp in length, we mapped fragments to a ~30 kbp region of the A1 genome (within that shown in Figure 3) and identified cases where the fragments spanned >70%. The ANI for each fragment is shown above the fragment in red and the ANI for the set of aligned fragments is shown to the right. Over the 8 examples, the average ANI is 91.8 ± 1.7%. The average length of aligned scaffolds to the ~30 kbp A1 genomic region is 27.2 ± 2.6 kbp (the sum of the aligned scaffolds is slightly longer in some cases). Mapping of reads to the scaffolds from Pig Lak 8 confirmed a high degree of within-population sequence variation that likely accounts for assembly fragmentation. **B.** Plot of ANI vs. fragment length for all (N=4830) pig fragments >2 kbp identified as possible Lak phage. We confirmed that the pig Lak genome fragments mapped throughout the A1 genome.

**Supplementary Figure 6.**
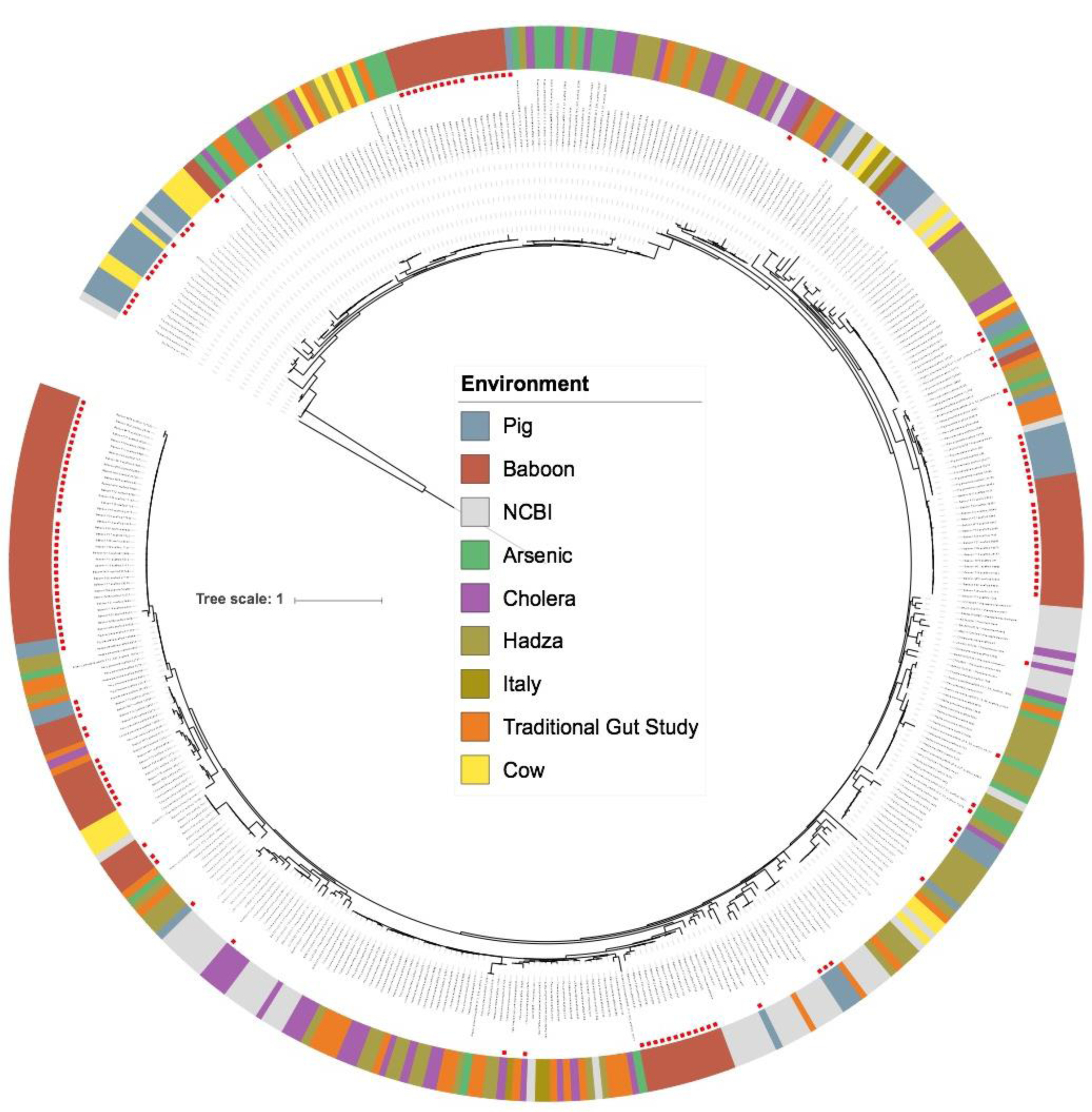
Phylogenetic tree of *Prevotella* from all cohorts examined in this study, constructed using 16S rRNA gene sequences on scaffolds >1 kbp in length, with information about the sample of origin. Red dots denote *Prevotella* strains that came from samples where Lak phage were also found. Note there is no strong separation of *Prevotella* type based on sample source and closely related strains come from samples with and without the megaphage. Data sources are as follows: Traditional gut study: (17), Baboon: (16), Hadza and Italy: (15), Cholera: (4), Cow: (21) and Pig data (this study). All data accession numbers listed in **Table S1**.

**Supplementary Figure 7.**
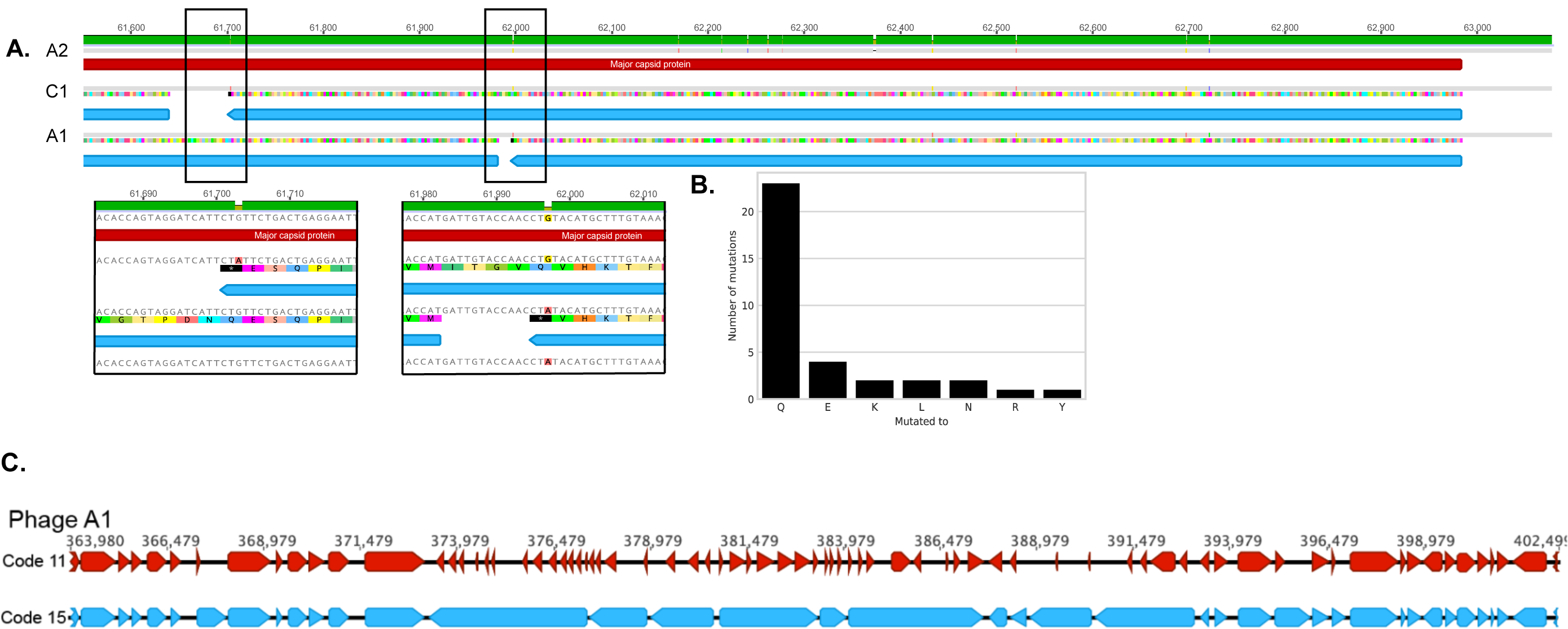
Evidence for alternative coding involving the stop codon TAG translated as glutamine (Q). **A**. An example of TAG mutated to an alternative codon encoding glutamine in A1, A2, and C1. **B**. Number of times across the three genomes that TAG is mutated to encode an amino acid. **C.** Gene predictions using genetic code 15 (blue open reading frames) *vs.* 11 (red open reading frames). Note some genes are not affected by the change from the prediction of TAG as a stop codon to encoding glutamine, whereas others are. Use of the alternative code corrected for clearly split genes and increased the predicted coding density from <70% to ~90%, genome wide.

**Supplementary Figure 8.**
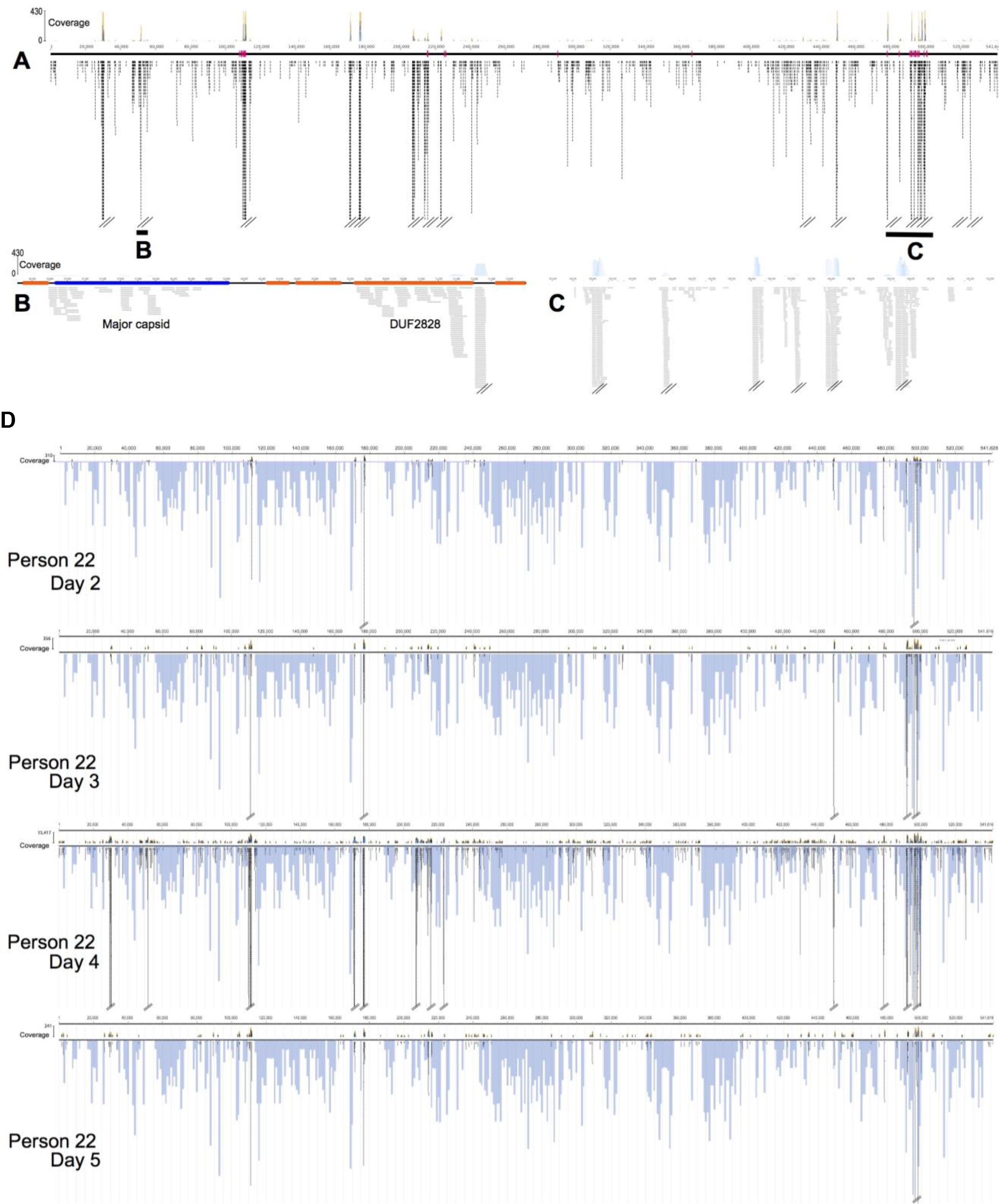
Mapping of transcript reads to the A1 megaphage genomes. **A.** Reads from 22-4. Red tick marks indicate the locations of tRNAs. Paired lines indicate read pile ups that have been truncated to save space (see coverage information in the top graph). Heavy black underlines indicate regions expanded in **B**. and **C**. Note the high level of expression of many intergenic regions, some expression of tRNAs and of certain proteins. **D**. Mapping of transcripts to the A1 genomes (as in A-C) superimposed on histograms showing the TAG use frequency in blue. In all cases, regions with and without the TAG codon are expressed.

**Supplementary Figure 9.**
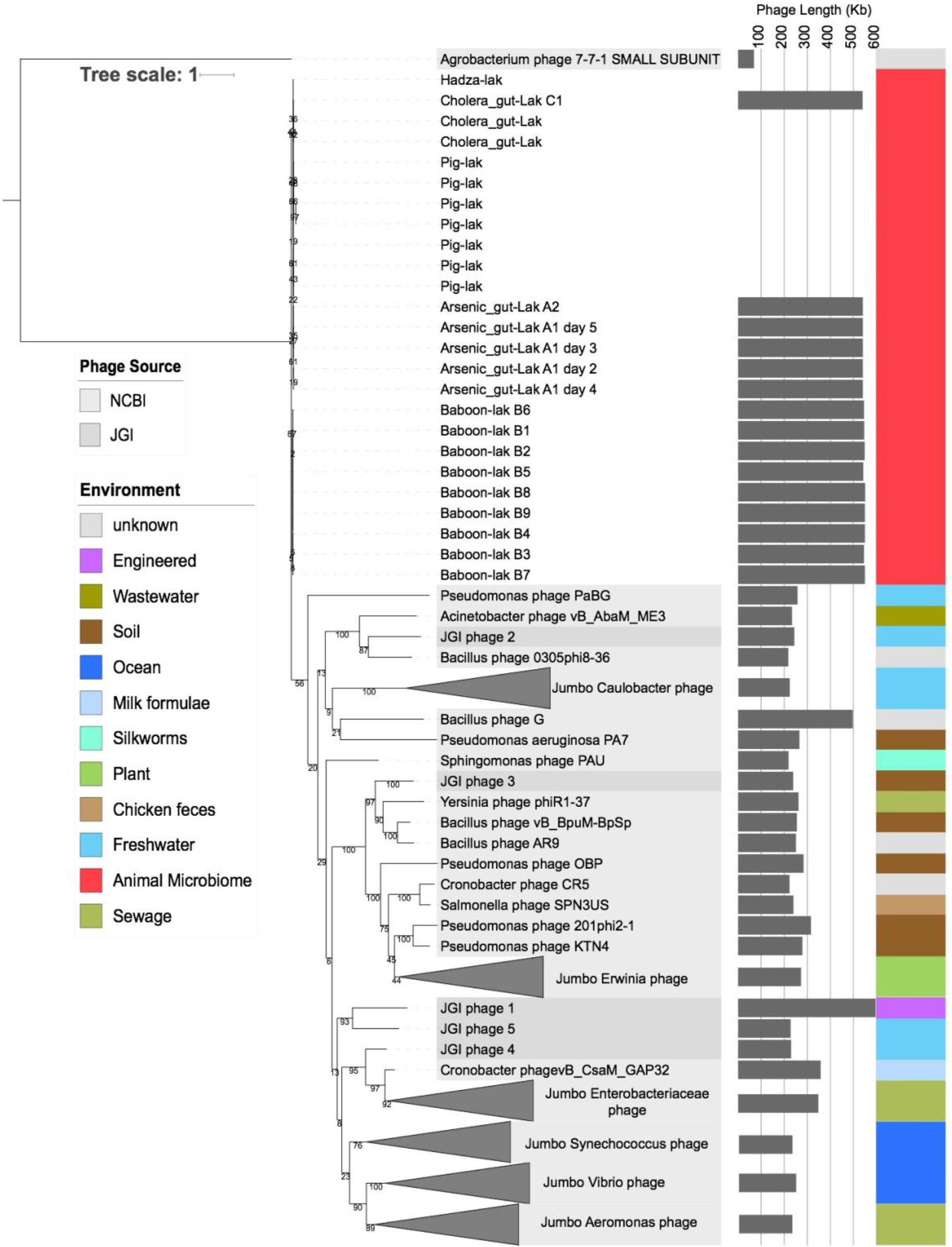
Phylogenetic analysis based on the terminase gene sequences from the genomes of the phage analyzed in this study and previously sequenced large phage genomes. Colored bars indicate the environment of origin. To our knowledge, the megaphage are the first phage genomes of >200 kbp in length recovered from a human body habitat. Grey bars indicate genome size (or average genome size for collapsed clades), and are not shown when the terminase was derived from an incomplete genome. Intriguingly, the terminase sequence is extremely well conserved in the megaphage reported here, despite significant genome divergence (**Table S3**). This unexpected sequence conservation mirrors that found in some tRNAs and tRNA introns.

**Supplementary Figure 10.**
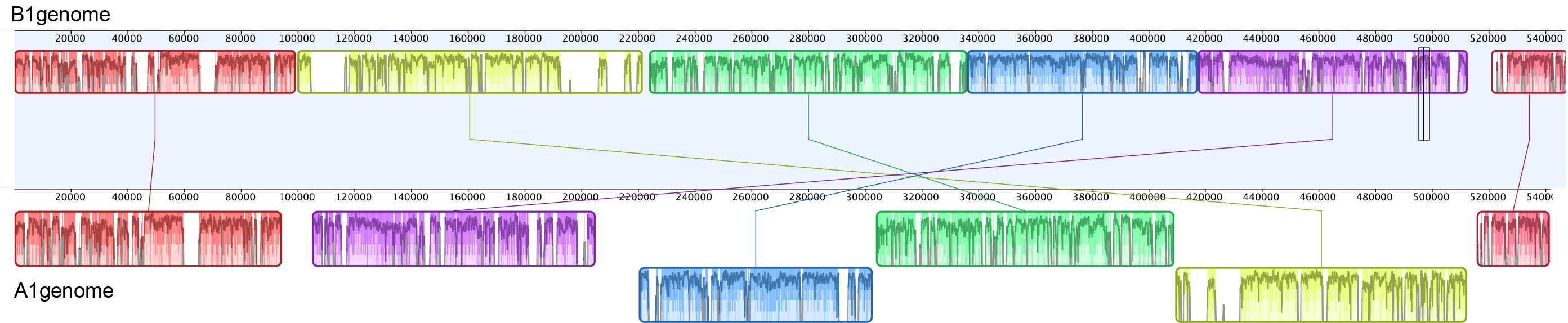
Diagram illustrating the rearrangements in the B1 relative to A-1, 2 and C1 genomes.

**Supplementary Figure 11.**
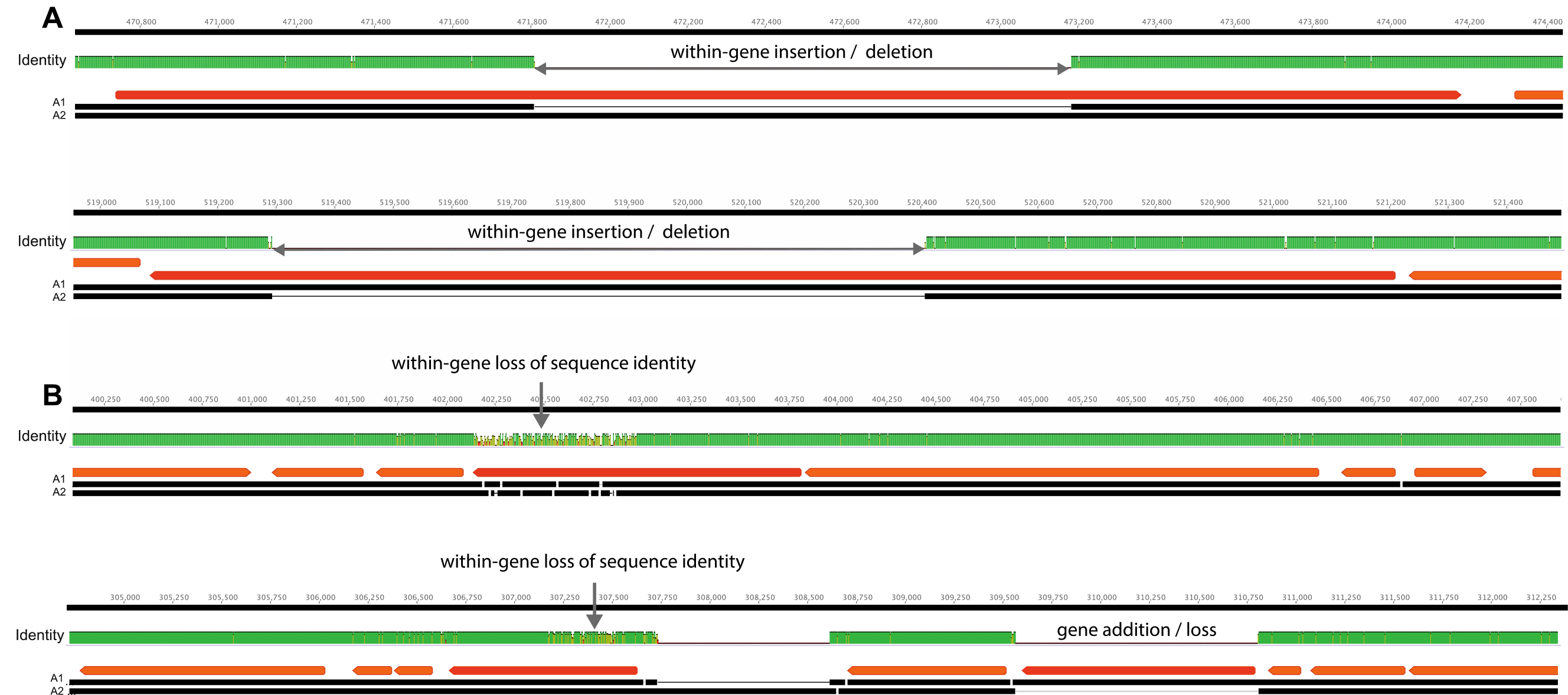
Examples of phenomena that distinguish the A1 and A2 genomes. **A.** Within-gene deletions, examples of which are also apparent in **Figure 3** alignments of all Lak genomes with A1. **B.** Sudden loss of sequence identity within a gene and an example of a gene insertion / loss that distinguishes the genomes.

**Supplementary Figure 12.**
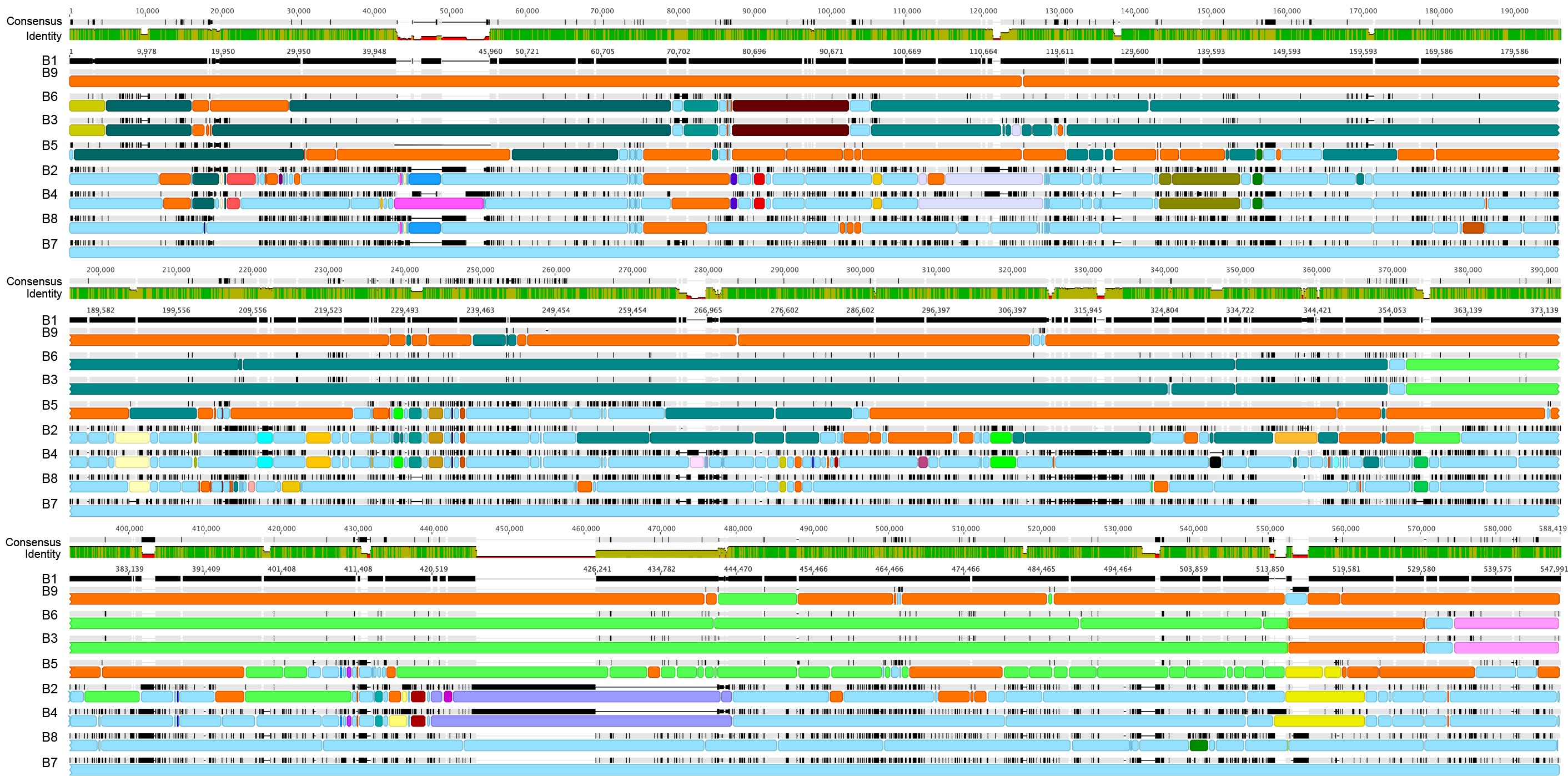
Alignment of the nine complete B-Lak genomes, with B1 (top black line) set as the reference sequence. Each genome is a row and vertical black tick marks indicate polymorphic sites. The sequence of B1 was designated as type ‘orange’ and sequence B7 as type ‘blue’. Colored underlines indicate sequence blocks that are identical. Small gaps within bars of the same color indicate SNPs that are not in agreement despite flanking sequence identity. Notably, the patterns indicate extensive admixture of sequence blocks, presumably due to homologous recombination, and often involve the available B-Lak genomes. For example, blocks of sequence shared with B7 (light blue underlines) occur in all B-Lak genomes. One region within the lowest panel is shown in detail in **Figure 4.**

**Supplementary Figure 13.**
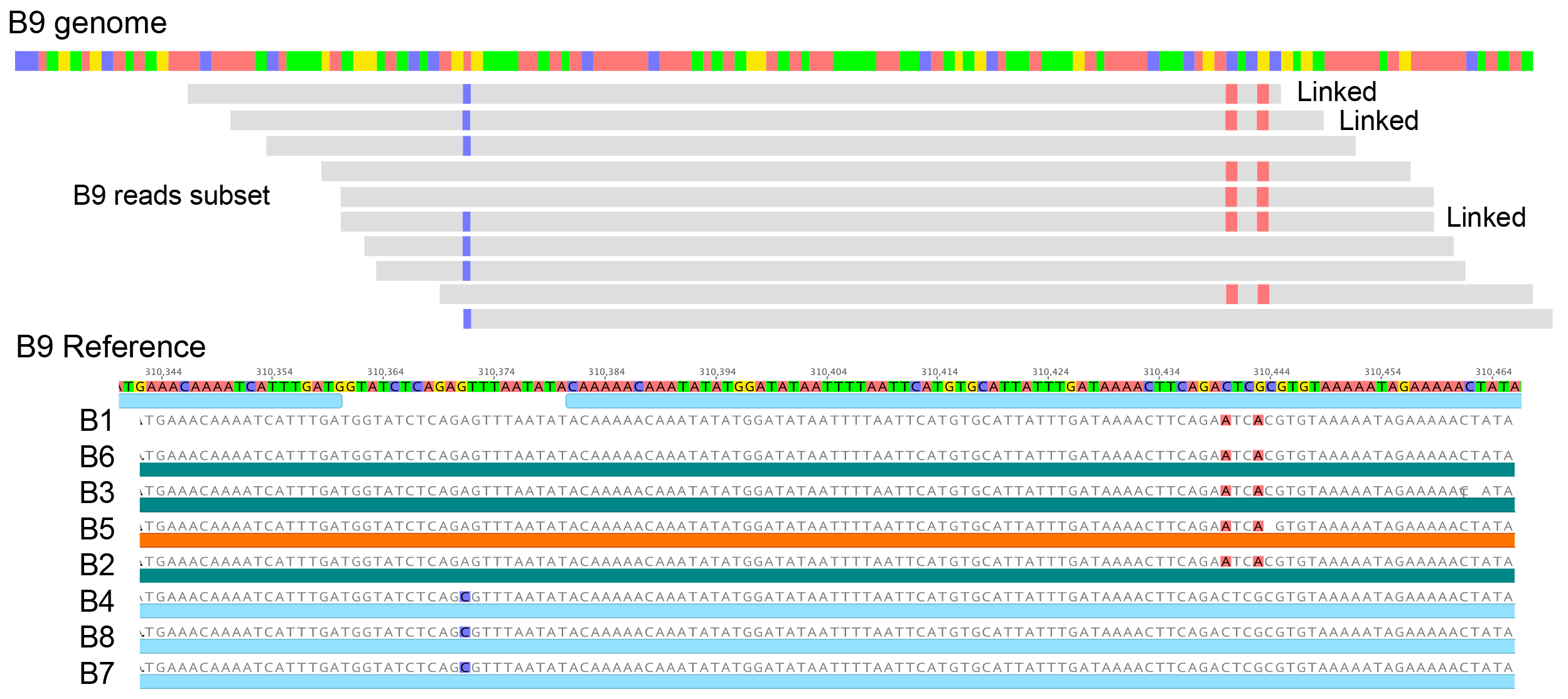
Diagram showing SNP-bearing single reads that span adjacent polymorphic sites and the comparison of these sequences to the B1-B9 genome sequences at this locus. Note that the blue and pair of red SNPs are not linked in the B1-B9 genomes but are linked in three sequencing reads. Thus, there is no single choice for the consensus sequence at this locus. The varied SNP linkage patterns are inferred to indicate homologous recombination.

**Supplementary Figure 14.**
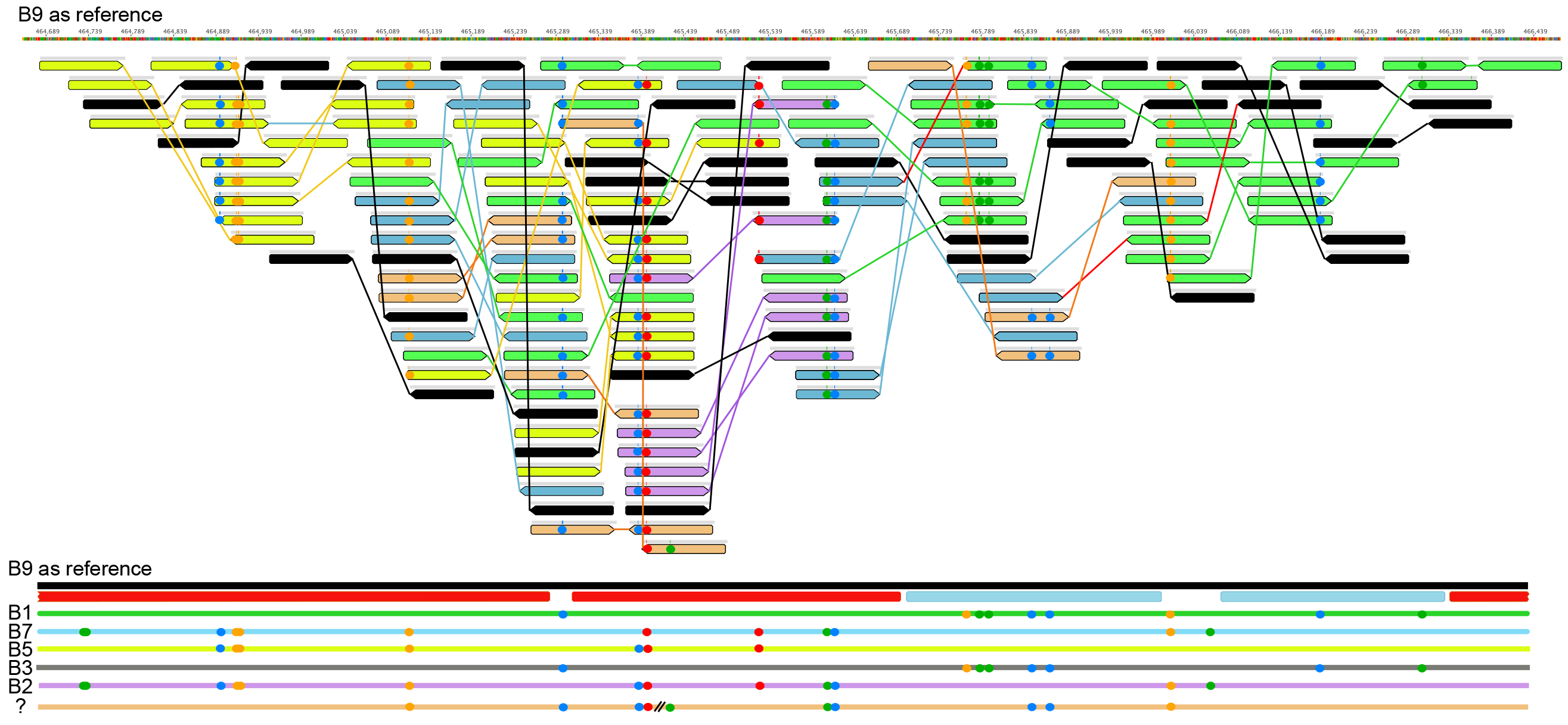
Genotypic variation within the B9 population. The upper panel shows a small subset of reads (light grey bars underlined with colored bars) that mapped to a hypervariable region of the B9 genome. Example paired reads were selected for display as they carried SNPs present in other reads (colored circles highlight SNP positions and types). As shown by red and blue bars in the lower panel, this region overlaps with that analyzed in **Figure 4**. Each read was assigned to a possible genotype (in some cases, multiple choices were possible), indicated by the color of the reads (top panel) so as to match to the color of the genome sequences in the lower panel. Despite relatively clonality of most other regions, these data indicate that the phage population includes rare genotypes that reflect homologous recombination, mostly involving sequences found in the B-Lak genome collection. Some paired reads contain combinations of SNPs not found in the B-Lak genomes (cream color). This may reflect recombination with other B-Lak genotype(s).

**Supplementary Figure 15.**
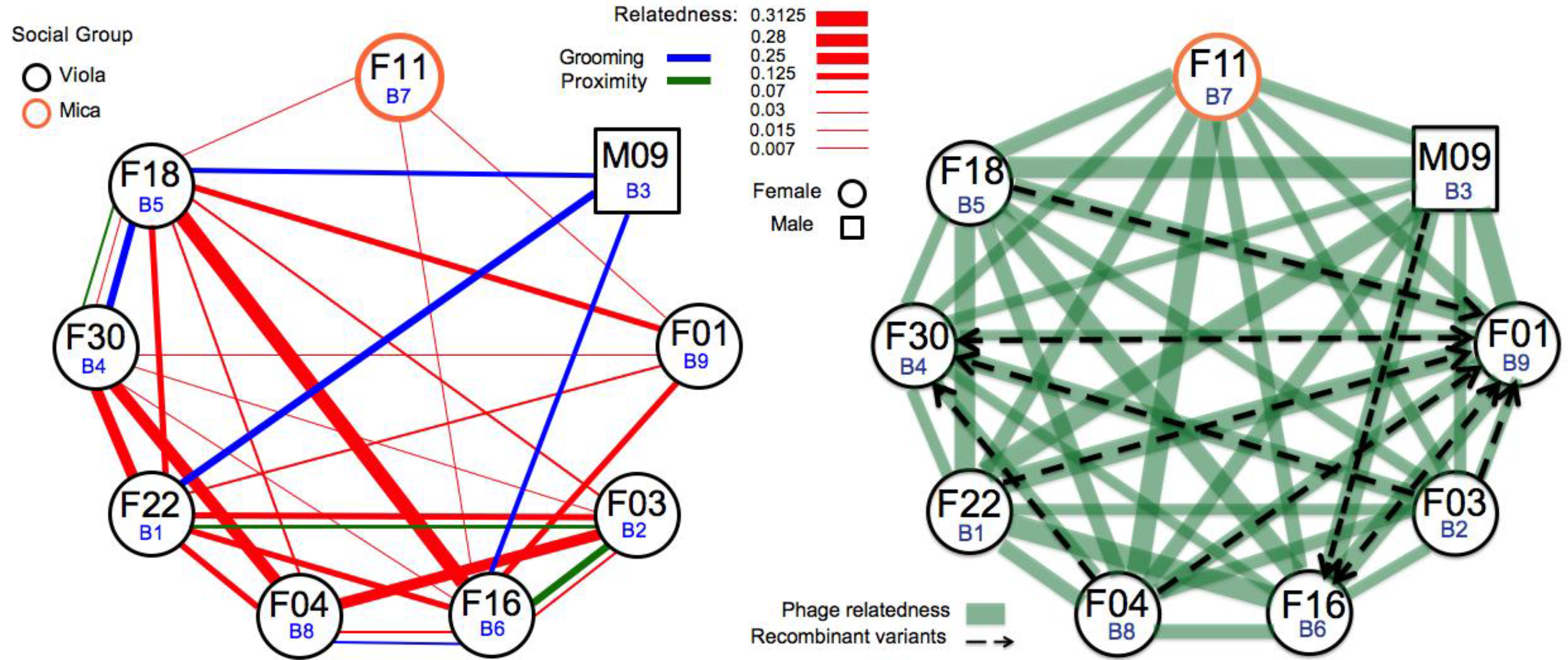
Summary of the metadata for the baboons and information about the relatedness of the phage populations. Left panel shows the baboon genetic relatedness (red lines) and social interactions (blue and green lines). All but one baboon comes from “Viola’s social group”. Right panel diagrams pairwise measures of phage genome sequence relatedness based on average nucleotide identity (ANI). Thicker lines indicate higher ANI. Recombinant variants refers to cases where more than about 0.1% of reads have SNPs that perfectly match sequences of another population (indicated by the arrow tip). In general, the baboon relatedness data do not predict phage relatedness or genotypic admixtures. For example, both F11 and M09 are essentially unrelated to the other baboons, as shown in the left panel, but their phage are about as strongly related to the phage of other baboons as are the phage in highly related baboons. Grooming interactions do not appear to lead to higher phage relatedness than occurs in the absence of these interactions. Similarly, proximity does not appear to be an important driver of phage relatedness (only strongest grooming and proximity links are shown but the full data are available in **Table S5**). Baboon metadata reported by (16).

**Supplementary Figure 16.**
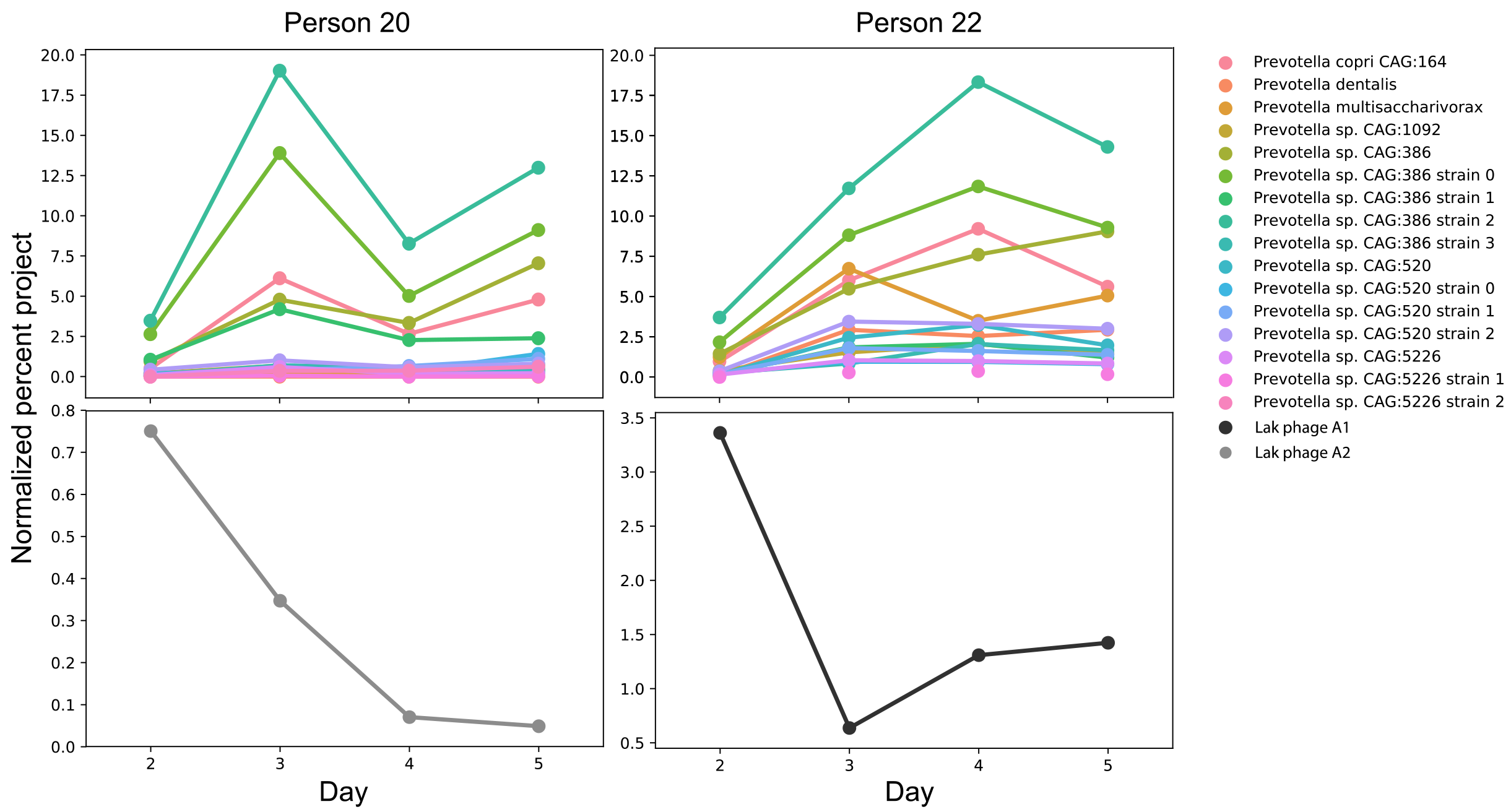
Diagrams showing the changing abundances of potential *Prevotella* hosts (top panel) and Lak phage (lower panel) over the four-days sampling period for Subject 20 and Subject 22 of the cohort of Laksam Upazila, Bangladeshi adults.

**Supplementary Figure 17.**
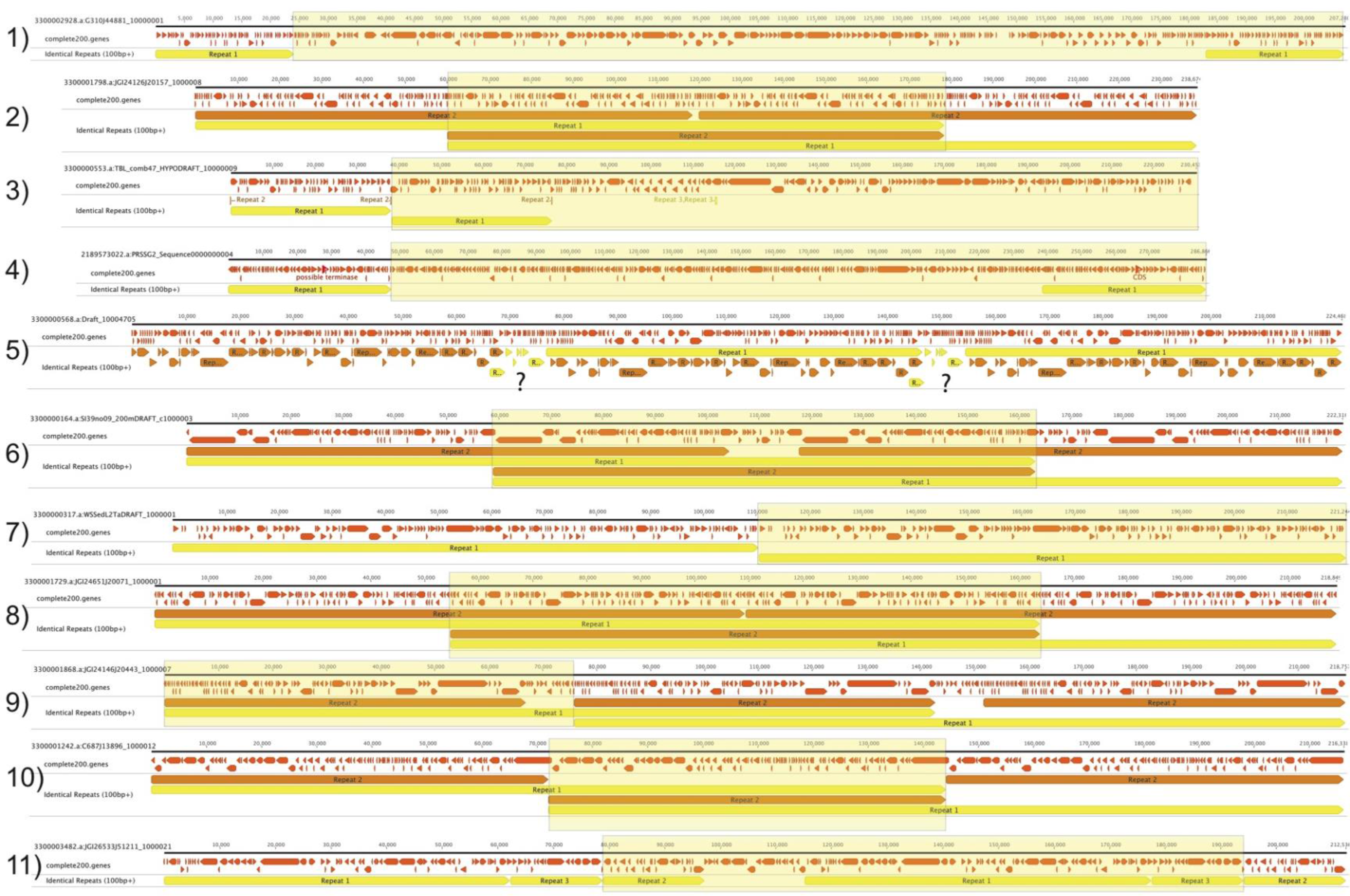
Analysis of questionable assemblies of reported >200 kbp phage genomes (14). Based on large perfect repeats, we suggest that these sequences are improbable. Gold highlights indicate our suggested alternative (and much shorter, in all but one case <200 kbp) chromosome choices.

## Supplementary Table Legends

**Supplementary Table 1**: Source of data used in this study

**Supplementary Table 2**: Comparison of tRNAs across the Lak megaphage genome set. **A**. The tRNAs in samples organized based on overall similarity in tRNA gene content. Note that some tRNA predictions are not very reliable (e.g., Sup tRNAs and pseudo-tRNAs) due to the large divergence between phage sequences and well studied tRNAs. **B.** The tRNAs in the Lak phage genomes listed based on position in the genome. Heavy black underlines indicate rearrangements to establish syntenic blocks

**Supplementary Table 3**: Table of average nucleotide identity (ANI) of phage genomes shared with A1, calculated over the aligned regions shown in **Figure 3B. B.** Lak phage ANI % over the full genome alignments.

**Supplementary Table 4**: The fraction of reads from each sample that map perfectly to the genome recovered from that sample and mapping of sequences carried on reads that do not map perfectly to other B-Lak genomes. Also shown is the fraction of reads that cannot be mapped to any reconstructed genome. An example mapping illustrates the case where unclassifiable SNPs tile out an otherwise unknown variant sequence. For more explanation, see **Supplementary Results**.

**Supplementary Table 5**: Baboon metadata. **A.** Baboon pedigree scores, **B.** Grooming interactions, **C**. Proximity scores. Data from (16). Based on **A**. and information in **Table S3**, genetic relatedness of baboons does not predict relatedness of phage, based on a correlation analysis.

**Supplementary Table 6**: Sequences and other information related to tRNA introns.

**Supplementary Table 7**: Genes with functional predictions. **A**. Listing of genes in the A1 Lak phage genome with a confident, moderately confident (purple functional prediction) or lower confidence (grey text) prediction. Structural proteins are listed in blue text. **B**. Genes with functional annotations from **Table S7A** sorted by function.

## REFERENCES

1. Sommer F, Bäckhed F (2013) The gut microbiota--masters of host development and physiology. Nat Rev Microbiol 11(4):227–238.

2. David LA, et al. (2014) Diet rapidly and reproducibly alters the human gut microbiome. Nature 505(7484):559–563.

3. Bäckhed F, et al. (2015) Dynamics and Stabilization of the Human Gut Microbiome during the First Year of Life. Cell Host Microbe 17(5):690–703.

4. David LA, et al. (2015) Gut microbial succession follows acute secretory diarrhea in humans. MBio 6(3):e00381–15.

5. Di Rienzi SC, et al. (2013) The human gut and groundwater harbor non-photosynthetic bacteria belonging to a new candidate phylum sibling to Cyanobacteria. Elife 2:e01102.

6. Waller AS, et al. (2014) Classification and quantification of bacteriophage taxa in human gut metagenomes. ISME J 8(7):1391–1402.

7. Yutin N, et al. (2018) Discovery of an expansive bacteriophage family that includes the most abundant viruses from the human gut. Nat Microbiol 3(1):38–46.

8. Minot S, et al. (2011) The human gut virome: inter-individual variation and dynamic response to diet. Genome Res 21(10):1616–1625.

9. Penadés JR, Chen J, Quiles-Puchalt N, Carpena N, Novick RP (2015) Bacteriophage-mediated spread of bacterial virulence genes. Curr Opin Microbiol 23:171–178.

10. Dutilh BE, et al. (2014) A highly abundant bacteriophage discovered in the unknown sequences of human faecal metagenomes. Nat Commun 5:4498.

11. Guerin E, et al. (2018) Biology and taxonomy of crAss-like bacteriophages, the most abundant virus in the human gut. bioRxiv:295642.

12. Gupta VK, Paul S, Dutta C (2017) Geography, Ethnicity or Subsistence-Specific Variations in Human Microbiome Composition and Diversity. Front Microbiol 8:1162.

13. Yuan Y, Gao M (2017) Jumbo Bacteriophages: An Overview. Front Microbiol 8:403.

14. Paez-Espino D, et al. (2016) Uncovering Earth’s virome. Nature 536(7617):425–430.

15. Rampelli S, et al. (2015) Metagenome Sequencing of the Hadza Hunter-Gatherer Gut Microbiota. Curr Biol 25(13):1682–1693.

16. Tung J, et al. (2015) Social networks predict gut microbiome composition in wild baboons. Elife 4. doi:10.7554/eLife.05224.

17. Obregon-Tito AJ, et al. (2015) Subsistence strategies in traditional societies distinguish gut microbiomes. Nat Commun 6:6505.

18. Andersson AF, Banfield JF (2008) Virus population dynamics and acquired virus resistance in natural microbial communities. Science 320(5879):1047–1050.

19. Hayashi H, Shibata K, Sakamoto M, Tomita S, Benno Y (2007) Prevotella copri sp. nov. and Prevotella stercorea sp. nov., isolated from human faeces. Int J Syst Evol Microbiol 57(Pt 5):941–946.

20. Ghosh TS, et al. (2014) Gut microbiomes of Indian children of varying nutritional status. PLoS One 9(4):e95547.

21. Thomas M, et al. (2017) Metagenomic characterization of the effect of feed additives on the gut microbiome and antibiotic resistome of feedlot cattle. Sci Rep 7(1):12257.

22. Das U, Shuman S (2013) Mechanism of RNA 2’,3′-cyclic phosphate end healing by T4 polynucleotide kinase-phosphatase. Nucleic Acids Res 41(1):355–365.

23. Serwer P, Hayes SJ, Thomas JA, Hardies SC (2007) Propagating the missing bacteriophages: a large bacteriophage in a new class. Virol J 4:21.

24. Mahmoudabadi G, Phillips R (2018) A comprehensive and quantitative exploration of thousands of viral genomes. Elife 7. doi:10.7554/eLife.31955.

25. Ambrozic J, Ferme D, Grabnar M, Ravnikar M, Avgustin G (2001) The bacteriophages of ruminal prevotellas. Folia Microbiol 46(1):37–39.

26. Ivanova NN, et al. (2014) Stop codon reassignments in the wild. Science 344(6186):909–913.

27. Denef VJ, Banfield JF (2012) In situ evolutionary rate measurements show ecological success of recently emerged bacterial hybrids. Science 336(6080):462–466.

28. Brown-Jaque M, Calero-Cáceres W, Muniesa M (2015) Transfer of antibiotic-resistance genes via phage-related mobile elements. Plasmid 79:1–7.

29. Castelle CJ, Banfield JF (2018) Major New Microbial Groups Expand Diversity and Alter our Understanding of the Tree of Life. Cell 172(6):1181–1197.

30. Hua J (2016) Capsid Structure and DNA Packing in Jumbo Bacteriophages. dissertation (University of Pittsburgh). Available at: http://d-scholarship.pitt.edu/27666/[Accessed June 19, 2018].

31. Sieber CMK, et al. (2017) Recovery of genomes from metagenomes via a dereplication, aggregation, and scoring strategy. bioRxiv:107789.

32. Langmead B, Salzberg SL (2012) Fast gapped-read alignment with Bowtie 2. Nat Methods 9(4):357–359.

33. Brown CT, et al. (2015) Unusual biology across a group comprising more than 15% of domain Bacteria. Nature 523(7559):208–211.

34. Kearse M, et al. (2012) Geneious Basic: an integrated and extendable desktop software platform for the organization and analysis of sequence data. Bioinformatics 28(12):1647–1649.

35. Hyatt D, et al. (2010) Prodigal: prokaryotic gene recognition and translation initiation site identification. BMC Bioinformatics 11:119.

36. Suzek BE, Huang H, McGarvey P, Mazumder R, Wu CH (2007) UniRef: comprehensive and non-redundant UniProt reference clusters. Bioinformatics 23(10):1282–1288.

37. Ogata H, et al. (1999) KEGG: Kyoto Encyclopedia of Genes and Genomes. Nucleic Acids Res 27(1):29–34.

38. Finn RD, Clements J, Eddy SR (2011) HMMER web server: interactive sequence similarity searchings. Nucleic Acids Res 39(Web Server issue):W29–37.

39. Zimmermann L, et al. (2018) A Completely Reimplemented MPI Bioinformatics Toolkit with a New HHpred Server at its Core. JMol Biol 430(15):2237–2243.

40. Lowe TM, Chan PP (2016) tRNAscan-SE On-line: integrating search and context for analysis of transfer RNA genes. Nucleic Acids Res 44(W1):W54–7.

41. Griffiths-Jones S, Bateman A, Marshall M, Khanna A, Eddy SR (2003) Rfam: an RNA family database. Nucleic Acids Res 31(1):439–441.

42. Biswas A, Staals RHJ, Morales SE, Fineran PC, Brown CM (2016) CRISPRDetect: A flexible algorithm to define CRISPR arrays. BMC Genomics 17:356.

43. Edgar RC (2010) Search and clustering orders of magnitude faster than BLAST. Bioinformatics 26(19):2460–2461.

44. DeSantis TZ, et al. (2006) Greengenes, a chimera-checked 16S rRNA gene database and workbench compatible with ARB. Appl Environ Microbiol 72(7):5069–5072.

45. Caporaso JG, et al. (2010) QIIME allows analysis of high-throughput community sequencing data. Nat Methods 7(5):335–336.

46. Edgar RC (2004) MUSCLE: multiple sequence alignment with high accuracy and high throughput. Nucleic Acids Res 32(5):1792–1797.

47. Stamatakis A (2014) RAxML version 8: a tool for phylogenetic analysis and post-analysis of large phylogenies. Bioinformatics 30(9):1312–1313.

48. Miller MA, Pfeiffer W, Schwartz T (2011) The CIPRES Science Gateway: A Community Resource for Phylogenetic Analyses. Proceedings of the 2011 TeraGrid Conference: Extreme Digital Discovery, TG ‘11. (ACM, New York, NY, USA), pp 41:1–41:8.

49. Letunic I, Bork P (2016) Interactive tree of life (iTOL) v3: an online tool for the display and annotation of phylogenetic and other trees. Nucleic Acids Res 44(W1):W242–5.

50. Darling ACE, Mau B, Blattner FR, Perna NT (2004) Mauve: multiple alignment of conserved genomic sequence with rearrangements. Genome Res 14(7):1394–1403.

51. Kurtz S, et al. (2004) Versatile and open software for comparing large genomes. Genome Biol 5(2):R12.

52. Bailly-Bechet M, Vergassola M, Rocha E (2007) Causes for the intriguing presence of tRNAs in phages. Genome Res 17(10):1486–1495.

